# Induced long-term potentiation improves synaptic stability and restores network function in ALS motor neurons

**DOI:** 10.1101/2025.05.04.652096

**Authors:** Anna M. Kollstrøm, Marthe Bendiksvoll Grønlie, Nicholas Christiansen, Axel Sandvig, Ioanna Sandvig

## Abstract

Amyotrophic lateral sclerosis (ALS) is a fatal neurodegenerative disease causing progressive dysfunction and degeneration of upper and lower motor neurons. An increasing body of evidence has identified synaptic alterations in patients and experimental models of ALS. Importantly, these have been associated with functional impairments in motor neuron networks, suggesting that synaptic impairments are early events in the disease cascade resulting in functional compensatory reconfigurations. The synapse may therefore represent a disease-modifying target to delay disease progression. In this study, we aimed to stabilize synapses and modify structural connectivity to restore network balance in ALS patient-derived motor neuron networks. To this end, we blocked the potassium channels using tetraethylammonium (TEA) which has been shown to induce chemical long-term potentiation (cLTP). The unperturbed ALS patient-derived motor neuron networks developed clear signs of subtle network dysfunction, including increased firing rate and bursting, and accompanying structural abnormalities. These features were partially restored by temporarily blocking the potassium channels. Specifically, the TEA-treated ALS networks were characterized by a reduction in aberrant branching and stabilization of dendritic spines, alongside a temporary reduction in firing rate and bursting. Furthermore, protein expression assays revealed restoration of dysregulated molecular pathways, including protein synthesis and metabolic pathways, and upregulation of pathways involved in synapse organization in the TEA-treated ALS networks. Collectively, these findings improve our understanding of the association between synaptic impairments and functional alterations in ALS, and demonstrate the relevance of modulating synaptic plasticity to promote network balance.

## Introduction

Despite an improved understanding of the pathophysiological mechanisms underlying neurodegenerative diseases, the failure to translate relevant findings into successful therapeutic outcomes remains a major challenge. A better understanding of the early-onset cellular and network responses to neurodegeneration is therefore urgently needed. Amyotrophic lateral sclerosis (ALS), the most common motor neuron disease, is a fatal neurodegenerative disease characterized by progressive motor impairment and paralysis, eventually resulting in respiratory failure (1). The most prominent hallmark of ALS is the degeneration of upper and lower motor neurons, however, there is growing evidence of early disruptions in neural circuit connectivity and activity, specifically increased functional connectivity in cortical and subcortical areas (2–4), and cortical hyperexcitability (5, 6). Widespread alterations in neuronal communication therefore appear to play a crucial role in disease progression before substantial cell death and neurological impairment are observed (7).

The maintenance of efficient neural network communication relies heavily on regulation of intrinsic neuronal firing mechanisms (8), balanced inhibitory and excitatory inputs (9, 10) and stable, but adaptable, synapses (11, 12). Structural and functional synaptic alterations have been observed in both ALS patients and animal models of the disease, resulting in disrupted neuronal signaling (13). Reduction of short interval-intracortical inhibition (SICI) has been identified in ALS patients using transcranial magnetic stimulation (TMS), indicating a loss of intracortical GABAergic circuits alongside increased activity of excitatory circuits (5, 14–16). Furthermore, findings from a SOD1-G93A mouse model of ALS have shown a temporary upregulation of inhibitory synapses in the spinal cord in early stages of ALS, followed by a subsequent breakdown of these inhibitory circuits (17). A selective loss of inhibitory interneurons innervating fast motor neurons has also been observed prior to motor neuron death (18), suggesting that early changes in inhibitory synapses contribute to network disintegration and disease progression.

Increased excitatory synaptic input and alterations in dendritic spines (19), changes in synaptic properties of the neuromuscular junction (NMJ) (20) and selective loss of tripartite synapses (21) have also been observed in presymptomatic and early stages of ALS. Importantly, dysregulated molecular pathways affecting the synapse have been identified in ALS patient-derived neurons and post-mortem patient tissue (22–24). Additionally, we have previously shown that disruptions in synaptic stability emerge before the onset of functional disturbances (25), indicating intrinsic synaptic vulnerability as an underlying cause of structural and functional network reconfigurations in ALS.

Synaptic impairment as an early event contributing to disease progression is further supported by studies aiming to restore synaptic function in ALS. Both motor behavior and motor neuron survival have been reported to improve by stabilizing spinal inhibitory synapses in mice (26), and restoration of synapses has been shown to increase neuronal survival and restore abnormal motor neuron firing (27, 28), underscoring the relevance of the synapse as a disease-modifying target. Crucially, to what extent synaptic alterations and the corresponding network reconfigurations represent pathogenic features or protective compensatory mechanisms that become maladaptive over time, remains unresolved (29, 30). Advanced engineered neural networks represent an avenue for targeted manipulation of vulnerable populations of neurons in early stages of ALS. These models recapitulate fundamental principles of brain network self-organization, including synaptic plasticity and emergent activity, as well as hallmarks of pathology (31–36). As such, they present an avenue for investigating the short- and long-term responses to early interventions on progressive neurodegeneration.

In this study, we aimed to enhance the synaptic strength and stabilize the synapses in ALS patient-derived motor neuron networks. By experimentally blocking potassium channels using tetraethylammonium (TEA), we manipulated the fundamental mechanism of neuronal excitability and action potential dynamics to engage critical aspects involved in the induction and expression of long-term potentiation (LTP) (37–40). Longitudinal electrophysiological and structural assays revealed that ALS networks were characterized by an increase in firing rate and bursting and showed signs of a more interconnected but less specialized network organization. This was accompanied by neurite outgrowth and branching abnormalities and a time-dependent loss of dendritic spines. Blocking of potassium channels led to temporary restoration of structural and functional disturbances to levels comparable to those of healthy controls. Furthermore, proteomic analysis of motor neuron synaptosomes revealed the restoration of ALS-associated dysregulated molecular pathways, including protein synthesis and metabolic pathways, and upregulation of pathways involved in synaptic plasticity in TEA-treated ALS networks. Together, our findings clearly demonstrate an association between synaptic dysfunction and disruptions in network activity in ALS and contribute to elucidate the role of synaptic plasticity in promoting neuroprotective mechanisms.

## Methods

### Human iPSCs and motor neuron differentiation

Human induced pluripotent stem cells (iPSCs) from a confirmed ALS patient with a C9orf72 mutation (female, 64; ID ND50001, NINDS Human Cell and Data Repository) and a healthy control donor (female, 49; ID ND50004, NINDS Human Cell and Data Repository) were expanded on vitronectin-coated (A14700, Thermo Fisher) 100 mm petri dishes using complete mTeSR Plus medium (mTeSR Plus Basal medium and mTeSR Plus 5X supplement; 100-0276, StemCell Technologies). The medium was exchanged daily. When the iPSCs reached approximately 80% confluence, motor neuron differentiation was initiated following the section “Procedure E” in the protocol published by Nijssen et al. (41) with minor modifications as previously described (25).

### Establishing networks on MEAs

Motor neurons were seeded on 6-well CytoView micro-electrode array (MEA) plates (M384-tMEA-6B; Axion Biosystems) on differentiation day 10. Two days before motor neuron seeding, MEAs were precoated with 0.05% poly (ethyleneimine) solution (PEI; P3141, Merck) diluted in HEPES (15630-056, Thermo Fisher) overnight. The PEI was removed the next day and the wells were rinsed four times with distilled water, before air drying the plates overnight. The next day, natural mouse Laminin (23017015, Thermo Fisher) was diluted in PBS (D8537, Sigma-Aldrich) to a final concentration of 20 *µ*g/mL, and the electrode area of the wells was coated for minimum 1 hour prior to seeding. Subsequently, the Laminin was removed and motor neurons were plated within the electrode area of each well at a density of 1500 neurons/mm^2^, yielding approximately 155,000 motor neurons per well. The cells were left to attach for 1 hour in the incubator (37°C, 5% CO_2_) before adding the final volume of medium, resulting in a total of 1 mL per well. Full media changes were performed on day 11, 12 and 13 of the differentiation protocol (41).

Human astrocytes at P8 (K1884, Thermo Fisher Scientific Gibco) were plated in the motor neuron networks at day 13. The astrocytes were thawed and centrifuged according to the manufacturer’s instructions, and resuspended in day 13 motor neuron medium. 90% of the media was replaced with fresh media in all wells, and the astrocytes were subsequently plated drop-wise in the remaining amount of media. The number of plated astrocytes corresponded to 10% of the motor neuron population. The networks were maintained with full media changes every other day for the remainder of the experiment.

### Chemical LTP induction

Chemical LTP (cLTP) was induced using the potassium channel blocker TEA, based on the protocol by Aniksztejn and Ben-Ari (37). 10% of the motor neuron media was removed from each well and replaced with 10% fresh media containing TEA to a final well concentration of 25 mM for 10 minutes. Half of the ALS networks received TEA. The remaining ALS networks and the healthy control networks only received media. After 10 minutes, all the TEA- or control media was removed and replaced with fresh media. The neural networks on 64-electrode MEAs were subsequently recorded for 59 minutes. cLTP was induced at three time points during the third week after seeding the neurons, at 27 days in vitro (DIV), 29 DIV and 31 DIV. Networks used for immunocytochemistry and mass spectrometry assays were fixed and collected immediately after removal of the TEA-containing media at 31 DIV. A timeline illustrating the full experiment is shown in Supplementary Fig. S1.

### Electrophysiological recordings

Neural network activity was recorded using the Axion Maestro Pro MEA system (Axion BioSystems, GA, USA). The AxIS Navigator software version 3.10.3 was used for data acquisition, sampled at 12.5 kHz. The plates were left for 10 minutes to allow network activity to stabilize after being moved to the recording platform, before starting a recording. Each recording lasted for 30 minutes. Recordings were made every other day from 26 DIV to 46 DIV, resulting in 11 recordings in total. Additionally, the networks were recorded on the same days as cLTP induction, as described in the section “Chemical LTP induction”. Spike detection was done with the AxIS Spike Detector, using an adaptive threshold of 7 standard deviations of the continuous data stream within frequency band 200Hz-3kHz.

### Immunocytochemistry and imaging

Motor neurons and astrocytes grown on 8-well chamber slides with removable media chambers (177402, Thermo Fisher) were used for immunocytochemistry assays. 50,000 motor neurons and 5000 astrocytes were plated in each well. Cells were fixed at 31 DIV, 38 DIV and 46 DIV with 3% glyoxal solution (42), containing 71% MQ water, 20% ethanol absolute, 8% glyoxal (128465, Sigma-Aldrich) and 1% acetic acid (1.00063, Sigma-Aldrich). The glyoxal was removed after 15 minutes and the samples washed three times with PBS (D8662, Sigma-Aldrich) for five minutes each round. 0.5% Triton-X (1.08643, Sigma-Aldrich) diluted in PBS was subsequently added to each well for five minutes to permeabilize the cells, before washing the samples two times with PBS for five minutes. A blocking solution made of 5% goat serum (G9023, Sigma-Aldrich) diluted in PBS was then added for 1 hour, leaving the samples in room temperature on an orbital shaker at 30 rpm. The blocking solution was replaced with PBS containing 5% goat serum and the primary antibodies: rabbit anti-Islet 1 (ISL1; 1:250; EP4182, Abcam), rabbit anti-HB9 (1:200; ab221884, Abcam), rabbit anti-ChAT (1:500; ab178850, Abcam), rabbit anti-Drebrin (1:100; PA5100142, Fisher Scientific), rabbit anti-GABA (1:100; A2052, Sigma-Aldrich), mouse anti-NeuN (1:1000; ab279295, Abcam), mouse anti-GluN1 (1:100; SAB5200546, Sigma-Aldrich), mouse anti-GluR1 (1:100; MA5-27694, Thermo Fisher), chicken anti-Neurofilament Heavy (NFH; 1:1000; ab4680, Abcam), chicken anti-GFAP (1:500; ab254083, Abcam), and chicken anti-MAP2 (1:5000; ab5392, Abcam). The samples were left with the primary antibodies on a tilting shaker in a cold room (3°C) overnight. The next day, the primary antibodies were removed and the samples washed three times with PBS, before incubating with the secondary antibodies in room temperature for three hours on an orbital shaker at 30 rpm. The following secondary antibodies were used: goat anti-rabbit Alexa Fluor 488 (0.2%; A11008, Thermo Fisher), goat anti-rabbit Alexa Fluor 568 (0.1%; A11011, Thermo Fisher), goat anti-mouse Alexa Fluor 647 (0.1%; A21236, Thermo Fisher), goat anti-chicken Alexa Fluor 568 (0.1%; ab175477, Abcam), and goat anti-chicken Alexa Fluor 488 (0.1%; A11039, Thermo Fisher). After removing the secondary antibodies, Hoechst diluted 1:2000 in PBS (bisbenzimide H 33342 trihydrochloride; 14533, Sigma-Aldrich) was added for 10 minutes to label nuclei, and the samples were subsequently washed three times with PBS for five minutes, before washing once with distilled water. The media chambers were removed and glass cover slides were mounted using anti-fade fluorescence mounting medium (ab104135, abcam). The samples were imaged using an EVOS M7000 microscope (Invitrogen) with an EVOS LWD 20x/0.45 NA (AMEP4982) objective, and EVOS M5000 microscope (Invitrogen) with an Olympus UPLSAPO, 20x/0.75 NA (N1480500) objective, and the following LED light cubes: DAPI (AMEP4650), GFP (AMEP4651), TX-Red (AMEP4655) and CY5 (AMEP4656). Image processing was performed in Fiji/ImageJ v1.54f.

### Synaptosome isolation

For synaptosome isolation, motor neurons and astrocytes were maintained on 6-well plates in parallel with MEAs and chamber slides. Synaptosomes were isolated immediately after the third cLTP induction at 31 DIV, using Syn-PER Synaptic Protein Extraction Reagent (87793, Thermo Fisher) and EDTA-free Halt Protease and Phosphatase Inhibitor Cocktail (78441, Thermo Fisher) according to the manufacturer’s instructions. The cells were washed twice using ice cold PBS, before adding Syn-PER and scraping off the cells. The lysate was transferred to a micocentrifuge tube and centrifuged at 1200 x *g* for 15 min. The supernatant was transferred to a new tube and centrifuged at 15,000 x *g* for 30 min. The supernatant was removed before the synaptosome pellet was resuspended in Syn-PER with protease and phosphatase inhibitor and DMSO, and kept in −80°C until analysis. All reagents were applied cold and the centrifugation steps were performed at 4°C.

### Mass spectrometry

Mass spectrometry (MS) analysis was performed by the Proteomics and Modomics Experimental Core Facility (PROMEC), Norwegian University of Science and Technology (NTNU). The synaptosome samples were resuspended in 100 *µ*L 1% sodium deoxycholate, 100 mM Tris-hydrochloride (pH 8.5), 10 mM Tris(2-carboxyethyl)phosphine, and 40 mM Chloroacetamide. The samples were sonicated 10 cycles of 30 s on and 30 s off, using the Bioruptor Pico sonicator (B01080010, Hologic Diagenode). Next, the samples were heated for 30 minutes at 95°C, before adding 100 *µ*L 0.01 M ammonium bicarbonate and 0.5 *µ*g trypsin for digestion overnight at 37°C. Peptides were desalted using C18 spin columns, and dried in a SpeedVac vacuum concentrator centrifuge (Thermo Fisher). The peptides were resuspended in 0.1% formic acid before performing liquid chromatography with tandem mass spectrometry (LC-MS/MS) using a timsTOF Pro with the nanoE-lute liquid chromatography system (both from Bruker Daltonics). The peptides were separated for 100 minutes using a 150 *µ*m*25 cm Pepsep 25 (Bruker Daltonics) with running buffers (A) 0.1% formic acid and (B) 0.1% formic acid in acetonitrile with a gradient of 2 - 40%. The timsTOF Pro was operated in the DDA PASEF mode with the following settings: 10 PASEF scans/cycle; accumulation and ramp times = 100 ms; target value = 20,000; dynamic exclusion activated = 0.4 min; quadruple isolation width = 2 Th for m/z < 700 and 3 Th for m/z > 800.

### Bioinformatic analyses and visualization

The raw MS files were analyzed using MaxQuant (version 2.4.14.0; (43)). MaxQuant output files were analyzed in RStudio (RStudio IDE 2024.09.1+394; (44)). Differentially expressed proteins (DEPs) were identified using the R package DEP (version 1.26.0; (45)). Proteins with missing values in more than one condition were excluded from further analysis. The data was normalized by variance stabilizing normalization, and the remaining missing values were imputed with the k-nearest neighbor approach for values missing at random. Proteins passing the thresholds Log2 fold change *≥* 1 and alpha=0.05 were considered as significantly differentially ex-pressed. However, the full list of DEPs were used for functional enrichment analyses to explore overall trends in the data and coordinated changes caused by groups of proteins (46). Functional enrichment analyses were performed using clusterProfiler (version 4.12.6; (47–50)). Kyoto Encyclopedia of Genes and Genomes (KEGG) pathway gene set enrichment analysis (GSEA) was performed with *Homo Sapiens* as organism and minimum gene set size set to 100. Gene Ontology (GO) over-representation analysis was performed on lists of upregulated and downregulated DEPs with Benjamini-Hochberg adjusted p-value<0.05, and plotted using enrichplot (51). GO GSEA was performed with number of permutations set to 10,000, minimum and maximum gene set sizes of 100 and 500, respectively, and Benjamini-Hochberg adjusted p-value<0.05. The SynGO geneset tool was used to identify overrepresented synaptic terms (52). Cytoscape was used to visualize networks of the overrepresented synaptic parent and child terms identified with the SynGO tool (53). Volcano plots were generated using EnhancedVolcano (54) with thresholds Log2 fold change *≥* 0.5 and Benjamini-Hochberg adjusted p-value<0.05. Plots were adjusted using ggplot2 (55). Information about individual proteins and their function was investigated using the website GeneCards: The Human Gene Database (56).

### Neurite network quantification

The Fiji/ImageJ (v1.54f) macro set NeuroConnectivity (57, 58) was used for quantification of neurite length, density and branching in neural networks immunolabeled with NFH and Hoechst. The networks were imaged using an EVOS M7000 microscope (Invitrogen) with an EVOS LWD 20x/0.45 NA (AMEP4982) objective, and the LED light cubes DAPI (AMEP4650) and TX-Red (AMEP4655). An automated scan protocol capturing 20% of the well area in a random order was used for image acquisition, resulting in 72 frames per well. The auto-focus setting was used at each capture and pixel sizes were 0.309 *µ*m x 0.309 *µ*m. The networks included in the analysis were fixed at two different time points, the first immediately after the final application of TEA at 31 DIV and the second one week later at 38 DIV. Four individual wells were imaged per group at each time point. The images were manually assessed after acquisition to exclude images with artifacts that could be incorrectly identified as neurites (i.e., images containing an edge of a well or large debris, or images out of focus). This resulted in the following number of images per group: Healthy 31 DIV, n = 262; Healthy 38 DIV, n = 236; ALS 31 DIV, n = 261; ALS 38 DIV, n = 242; ALS (+TEA) 31 DIV, n = 267; ALS (+TEA) 38 DIV, n = 266. Neurite detection was based on (59) and the Stardist algorithm (60) was used for detecting nuclei.

### Dendritic spine quantification

Confocal imaging for dendritic spine quantification was performed by the Cellular & Molecular Imaging Core Facility (CMIC) at NTNU using a Nikon Crest X-Light V3 spinning disk confocal microscope. To identify regions with motor neurons, overview images were acquired in widefield mode at 4x and 10x magnification with only the MAP2 channel. Six regions were picked per well, capturing mainly dendrites rather than cell bodies. The regions were picked at low magnification and without the drebrin channel, and were distributed out for each area but avoiding regions too close to the edges or otherwise damaged regions. The six regions per well were then imaged at higher magnification with a spinning disk of 50 *µ*m pinhole size inserted, using a 100x oil immersion objective with numerical aperture 1.45 and with the emission iris at 30% opening. A z stack with 200 nm slice thickness was acquired for each region. A Photometrics Kinetix back-illuminated sCMOS camera was used in 16-bit sub-electron mode for the high magnification acquisition, with an exposure time of 163 ms for each channel. A Lumencor Celesta light engine laser was used for excitation, with laser powers 5% for 546nm, 8% for 477nm and 5% for the 405nm excitation. Single band pass emission filters were used for each channel, 595/31, 511/20 and 438/24, respectively. Each channel was acquired per z plane before changing to the next z plane, to ensure acquisition at identical z planes. Following these procedures, 24 z stacks were obtained per group for each of the time points. All raw images were denoised using the Nikon NIS Elements AR nis.ai function (version 6.10.01). Fiji/ImageJ v1.54g was used for maximum intensity projection. 0.35% saturation clipping was performed for the MAP2 and drebrin channels to enhance contrast before quantifying dendritic spines. For each image, 10 *µ*m long segments of three individual secondary or tertiary dendrites were selected for manual spine quantification by a blinded investigator. This provided a total of 432 dendritic segments, i.e., 72 segments per group per time point. Only segments with drebrin-positive points that partially or fully overlapped with MAP2 were included in the analysis.

### Data analysis and statistics

Electrophysiology data from the MEAs was processed using the Python programming language (Python Software Foundation) with the Pandas package, based on the analysis by Weir et al. (61). Statistical analysis of electrophysiology and image quantification data was performed using the packages Scipy (62), Pingouin (63) and Scikit-posthocs (64), and figures were generated using Seaborn and Matplotlib.

Electrodes with a firing rate below 0.2 Hz and wells with a well firing rate below 0.2 Hz were excluded from network analysis. The well firing rate threshold for the recordings immediately following the TEA perturbation was set to 0.1 Hz. An overview of the number of networks included in network activity analysis is shown in Supplementary Table S1. The firing rate *f* was defined as *f* = (*spikes −* 1)*/n*Δ*t*, where *spikes* is the total number of recorded spikes in a given well, *n* is the number of electrodes considered to be active throughout the recording and Δ*t* is the time difference between the first and the last recorded spike.

The instantaneous firing rate *f*_*window*_ was defined as *f*_*window*_ = *spikes/n*Δ*t. Spikes* is the total number of spikes in the given time window, *n* is the number of active electrodes and Δ*t* the width of the given window. The instantaneous firing rate was calculated as a moving window with window size of 100 ms and step size of 10 ms, resulting in partially overlapping windows.

The inter-spike interval (ISI) was defined as the time interval between two consecutive spikes. Bursts were defined as a train of at least four spikes with an ISI less than 100 ms on a single electrode. The burst frequency *f*_*burst*_ was defined as *f*_*burst*_ = *bursts/n*Δ*t*, with *bursts* being the total number of bursts throughout the recording in a given well, *n* is the number of active electrodes and Δ*t* the time difference between the first and the last recorded spike. The inter-burst interval (IBI) was defined as the time interval between two consecutive bursts. Based on these definitions, the mean duration of bursts and the fraction of spikes in bursts were identified. The coherence index was used to assess synchrony of network activity (65).

Network bursts were defined as a sequence of at least four spikes across a well with an ISI_4_ less than 100 ms, meaning that a sequence of at least four spikes with the time between the first and the last spike considered was lower than the set threshold as described in Bakkum et al. (66). The network burst was continued until the ISI_4_ was longer than the given threshold.

Functional connectivity was detected by identifying the co-occurrences of spikes between any given pair of electrodes in each well. Spikes were binned to time windows of 100 ms such that a bin was considered active if it contained at least one spike. The number of bins considered active at the same time for each pair of electrodes were counted and normalized by the largest co-occurence value for the given well. This yielded networks consisting of nodes corresponding to the electrodes and weighted edges from the normalized co-occurences for each electrode pair. The networks were pruned by iterating through all nodes and removing the weakest connection from each node starting with the most connected node. This process was repeated until the desired density was reached, starting again with the most connected node when all nodes had been pruned once. To avoid disconnected nodes in the networks, nodes with two or fewer connections were left as is. Networks which were already sparser than the desired threshold were not pruned. Pruning was performed to obtain networks of lower density, and reach a more uniform distribution of network densities across the three groups. A density threshold of 50% was selected for further analysis, as this ensured the most evenly distributed network densities. Supplementary Table S2 shows the number of networks per group included in the connectivity analysis at each DIV. The giant component was extracted, and various metrics from graph theory were determined using the python package NetworkX (67). Mean degree refers to the sum of edges connected to a given node. The mean clustering of a node describes how well-connected its neighbors are, i.e., to what extent its neighbors are connected to each other. Distance based metrics were calculated using the inverse weight, as strongly connected nodes are considered to be in closer proximity than nodes with weaker connection strengths (68). This included the average path length, network diameter, and the betweenness centrality. The path length represent the average shortest path, i.e., number of steps, between all possible pairs of nodes. The diameter of the network is defined as the longest of the shortest paths between all pairs of nodes. The betweenness centrality provides an estimate for how often a node lies on the shortest path of other node pairs. The global efficiency describes how efficiently the network transfers information, and is defined as the inverse of the average shortest paths between the nodes. The local efficiency is the global efficiency of the subgraph comprised of the neighbors of a given node. The Small-World Propensity was calculated using the methods described in Muldoon et al. (68), and provides a measure for the extent to which the network displays characteristics of a small-world architecture, i.e., short average path length and high clustering.

The groups were assessed for normality at each individual time point using the Shapiro-Wilk’s test. Statistical comparisons between two groups were conducted using Student’s t-test. Comparisons between all three groups were conducted using One-Way ANOVA in the case of normally distributed data or the Kruskal-Wallis test in the case of non-normality, followed by Conover’s test for multiple comparisons with Bonferroni correction. Mixed-effects models were applied to investigate differences over time in the different network parameters between the ALS group, the TEA-treated ALS group and the healthy control group. The models were implemented in R using the lme4 package (version 1.1-35.5; (69)). Each network parameter was used as response variable and the interaction between the age of the networks (DIV) and condition (ALS, ALS (+TEA) or healthy control) was used as a fixed effect. Each well was used as a random effect to account for variation between each individual network structure. Model fit was evaluated based on the Akaike Information Criterion (AIC), and further evaluated through analysis of the model’s residuals, using the DHARMa package (70). When the data was normally distributed, a gaussian distribution with an identity link function was considered the best fit, and a linear mixed-effects model (LMM) was applied. In the case of non-normality, generalized linear mixed-effects models (GLMMs) were fitted. Supplementary Table S3 shows the chosen distributions and link functions for each network parameter. Post-hoc tests for multiple comparisons were conducted using the Estimated Marginal Means (emmeans package version 1.10.6; (71)) with Bonferroni correction. Results were determined as significant when p<0.05.

## Results

### Reduced structural network abnormalities and stabilization of dendritic spines indicate synaptic potentiation following blocking of potassium channels

Immunocytochemistry assays at 46 DIV confirmed the neuronal identity of both healthy control and ALS networks through the expression of NeuN and motor neuron markers ISL1, HB9 and ChAT (Supplementary Fig. S2). Further characterization also revealed positive expression of cytoskeletal markers MAP2 and NFH, the AMPA-receptor subunit GluR1, NMDA-receptor subunit GluN1, and the inhibitory neurotransmitter GABA. The presence of astrocytes was also confirmed with the expression of GFAP (Supplementary Fig. S2).

To investigate the effects of synaptic potentiation on network morphology, neurite density, length and branching were quantified in networks immunolabeled with NFH and Hoechst (Fig. 1A) on the same day as the final application of TEA (31 DIV) and one week later, at 38 DIV. The neurite density, represented by the total area covered by neurites, was higher in the untreated ALS networks compared to the healthy networks at both 31 DIV and 38 DIV (p<0.001; Fig. 1B). Interestingly, the neurite density was lower in the TEA-treated ALS networks compared to that of the untreated ALS networks (p<0.001), and showed a tendency toward the healthy networks, but was still significantly higher than the healthy networks (p<0.001). At 38 DIV, the neurite density remained lower in the TEA-treated ALS networks compared to the untreated ALS networks (p<0.001), and appeared comparable to the healthy networks although lower at this time point (p<0.001). The average neurite length was shorter in the untreated ALS networks compared to the healthy networks at 31 and 38 DIV ((p<0.001); Fig. 1C). The TEA-treated ALS networks again showed a trend similar to that of the healthy networks, and the average neurite length was longer compared to the untreated ALS networks at 31 DIV (p<0.001), but still shorter than the healthy networks (p<0.001). The neurite length remained shorter in both ALS groups compared to the healthy controls at 38 DIV (p<0.001), and no significant difference between the untreated and the TEA-treated ALS networks was observed at this time point (p=0.0963). The total number of neurite branches (Fig. 1D) revealed increased branching in the untreated and the TEA-treated ALS networks compared to the healthy networks at both 31 and 38 DIV (p<0.001). However, the TEA-treated ALS networks exhibited less branching compared to the untreated ALS networks (p<0.001), and appeared comparable to the healthy networks, especially at 38 DIV, yet not significantly so. Taken together, the neurite network quantification revealed reduced neurite length but increased density and branching in the ALS networks compared to the healthy networks, whereas these differences were partly modified and reduced following blocking of the potassium channels in ALS networks.

**Fig. 1.**
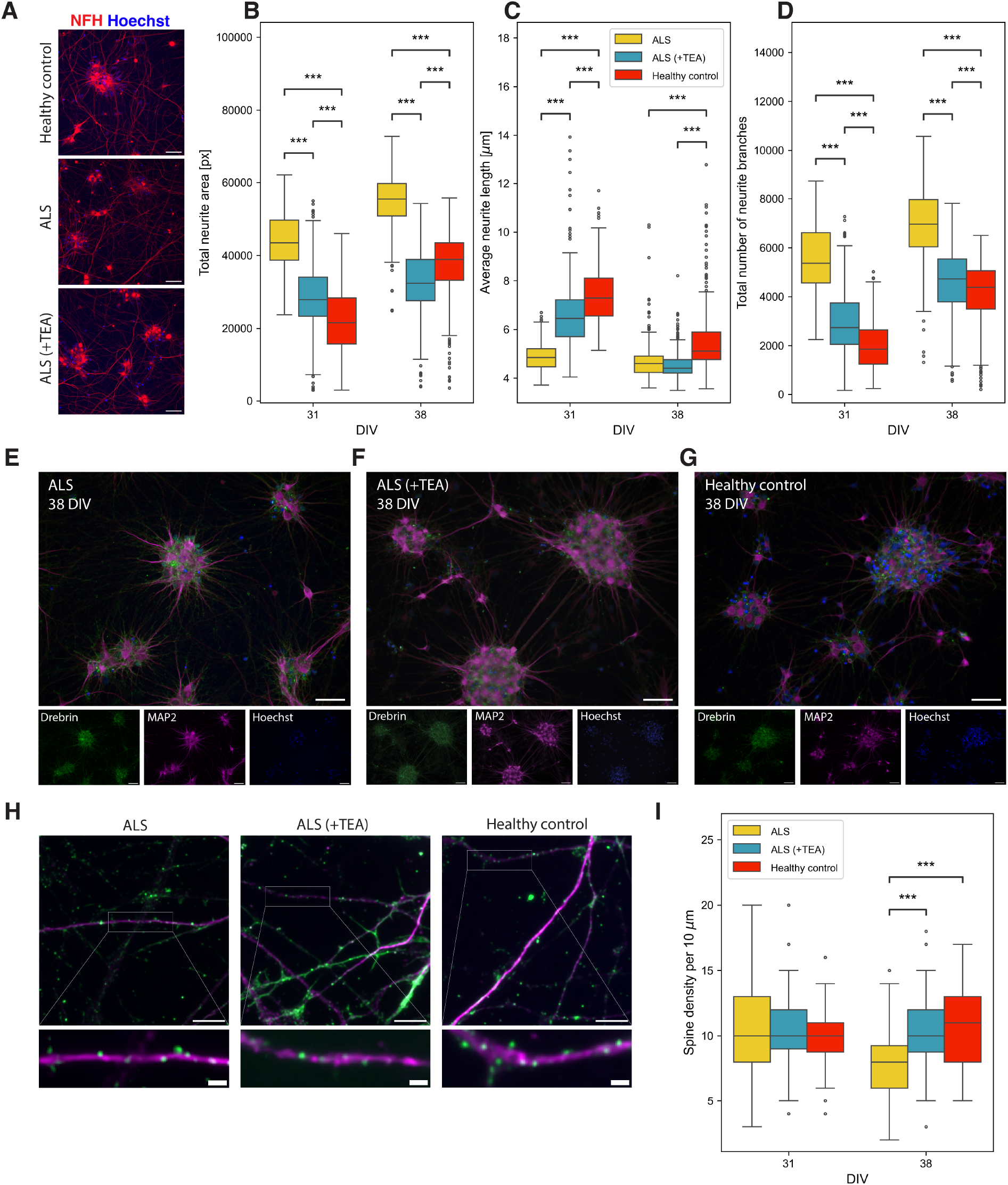
Network structure abnormalities are partly recovered after blocking of the potassium channels in ALS networks. (**A**) Example images of healthy, ALS and ALS (+TEA) motor neuron networks immunolabeled with Neurofilament Heavy (NFH). (**B-D**) Box plot showing group differences between ALS, ALS (+TEA) and healthy controls from the neurite length quantifications, with the total area covered by neurites (B), average neurite length (C) and total number of neurite branches (D). Scale bars: 100 *µ*m, legend shared between B-D. (**E-G**) Example images of ALS, ALS (+TEA) and healthy control motor neuron networks immunolabeled with drebrin and MAP2, and Hoechst for nuclei. Scale bars: 100 *µ*m. (**H**) Confocal images of clusters of drebrin along 10 *µ*m dendritic segments were used for quantification of dendritic spine density. Scale bars: 10 *µ*m (top) and 1 *µ*m (bottom). (**I**) Box plot showing group differences in spine density between ALS, ALS (+TEA) and healthy controls at 31 and 38 days in vitro (DIV). Kruskal-Wallis test, followed by Conover’s test for multiple comparisons with Bonferroni correction. *** p<0.001. TEA, tetraethylammonium.

Structurally, changes in synaptic strength are associated with dendritic spine remodeling (39, 72), and dendritic spine abnormalities have been associated with a number of neurodegenerative diseases, including ALS (19, 73). We therefore next assessed whether the blocking of potassium channels led to structural changes in spine density. Networks immunolabeled with the actin-binding protein drebrin and MAP2 were used for quantification (Fig. 1E-G). As drebrin is highly accumulated in dendritic spines, we estimated the density of drebrin clusters along 10 *µ*m dendritic segments (Fig. 1H). No differences in spine density were observed between the untreated ALS group, the TEA-treated ALS group and the healthy control group at 31 DIV (Fig. 1I). However, while a decrease in spine density was observed in the untreated ALS networks from 31 to 38 DIV (p<0.001), no indications of spine loss were observed over time in neither the healthy networks (p=0.210) nor the TEA-treated ALS networks (p=0.591). In fact, the spine density was significantly higher in these two groups compared to the untreated ALS group at 38 DIV (p<0.001), indicating a time-dependent loss of spines in the ALS networks, which was stabilized in networks receiving TEA, consistent with synaptic strengthening.

### Altered network activity in ALS motor neurons is mitigated by blocking potassium channels

Longitudinal MEA recordings of neural network activity revealed an overall higher firing rate in the untreated ALS networks compared to the healthy control networks (Fig. 2A). The TEA-treated ALS networks were not significantly different from the untreated ALS networks at 26 DIV, prior to the first TEA application (p=0.835; Student’s t-test). However, on the days of TEA application (27, 29 and 31 DIV) and the following day (28, 30 and 32 DIV) the TEA-treated ALS networks were more comparable to the healthy networks, with a lower firing rate (Fig. 2A) and higher mean ISI (Supplementary Fig. S3A-B) than the untreated ALS networks. Nevertheless, no significant differences between the three groups were observed at any of the perturbation time points. From 34 DIV onward, the firing rate of the TEA-treated ALS networks returned to similar levels as those of the untreated ALS networks.

**Fig. 2.**
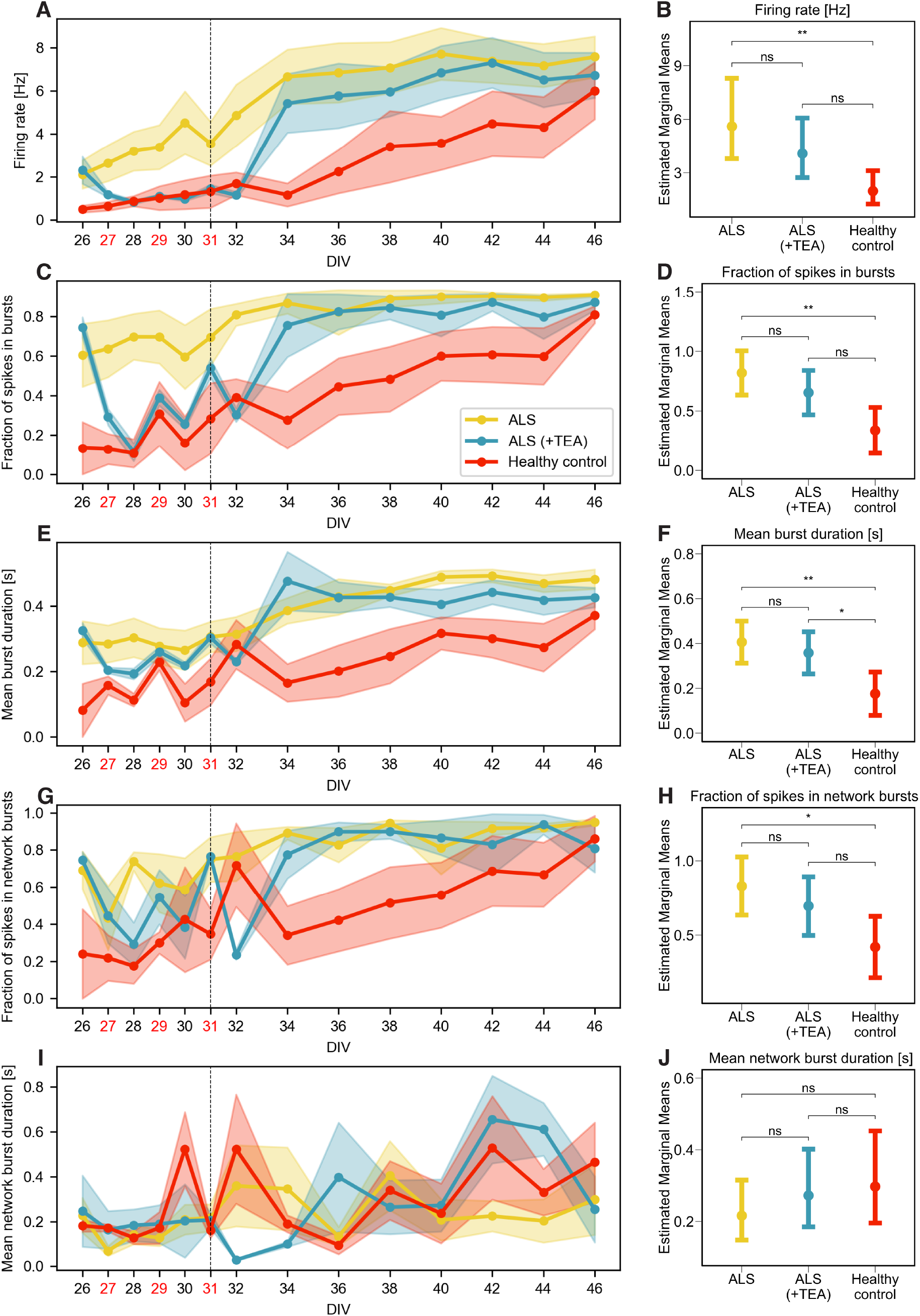
Differences in network activity relative to healthy control and untreated ALS networks before and after blocking of potassium channels. (**A-B**) Firing rate in Hz, (**C-D**) fraction of spikes in bursts, (**E-F**) mean burst duration in seconds, (**G-H**) fraction of spikes in network bursts, and (**I-J**) mean network burst duration in seconds. Left column plots: Longitudinal development of network activity. The mean values for all networks in each group are represented by the solid lines and circles. Shaded area shows the SEM. Tetraethylammonium (TEA) was applied to block potassium channels on the days labeled in red, that is 27, 29, and 31 days in vitro (DIV), and the vertical line at 31 DIV marks the final day of TEA application. Legend is shared between figures. Right column plots: Mixed-effects model estimated marginal means with 95% confidence interval for each of the metrics. * p<0.05, ** p<0.01, *** p<0.001. ns, not significant.

To investigate the network development over time, mixed-effects models were applied to statistically compare the groups across the recordings, as such models consider the dependence between repeated measurements. The three recordings immediately following the TEA application at 27, 29 and 31 DIV were not included in this analysis. An overview of all the mixed-effects models fitted and their estimated marginal means with 95% confidence interval is shown in Supplementary Table S3. This revealed a significant difference in firing rate between the ALS and the healthy networks over time (p<0.01), but no significant difference between the TEA-treated ALS networks and the untreated ALS networks (p=0.785), nor the healthy networks (p=0.0567; Fig. 2B). No significant differences between any of the groups were observed in the mean amplitude (Supplementary Fig. S3C-D). The coherence index, a measure of network synchrony, was transiently reduced in the TEA-treated ALS networks to the level of the healthy controls during and one day after perturbation, before increasing again and stabilizing at a similar level to that of the untreated ALS networks (Supplementary Fig. S3E). Overall however, no significant differences were observed in the coherence index between any of the groups (Supplementary Fig. S3F).

The untreated ALS networks were also characterized by increased bursting compared to the healthy networks, as evidenced by higher fraction of spikes in bursts (p<0.01; Fig. 2C-D) and longer burst duration (p<0.01; Fig. 2E-F). In contrast, the TEA-treated ALS networks followed a trend similar to the healthy networks, and were characterized by reduced burst frequency (Supplementary Fig. S4A-B) and burst duration on the days of the TEA application and one day after, but stabilized around the same levels as the untreated ALS networks from 34 DIV onward. Mixed-effects models revealed that the TEA-treated ALS networks were not significantly different from the untreated ALS networks nor the healthy networks in any of the burst parameters (Fig. 2C-F and Supplementary Fig. S4A-D), with the exception of the mean burst duration, which was lower in the healthy networks (p<0.05; Fig. 2F). Despite variations from time point to time point, the network burst metrics revealed higher network burst frequency (Supplementary Fig. S4E-F) and fraction of spikes in network bursts (Fig. 2G-H) in the ALS networks compared to their healthy counterparts (p<0.05). The TEA-treated ALS networks on the other hand, were not found to be significantly different from either of the other two groups, but showed a trend of reduced network burst frequency and fraction of spikes in network bursts compared to the untreated ALS networks. Interestingly, the mean network burst duration was slightly longer in the healthy and TEA-treated networks compared to the untreated ALS networks (Fig. 2I-J), albeit not significantly. Altogether, these findings identified several signs of subtle network dysfunction in the ALS networks, including higher firing rate, increased bursting and burst duration, and higher fraction of spikes in network bursts. All of these were temporarily reduced to a level comparable to the healthy controls following potassium channel blocking.

### Increased connectivity in ALS networks

Functional connectivity analyses were conducted to further investigate the information processing and computational capacity of the motor neuron networks. The connection strength increased over time in all three groups and followed a heavy-tailed distribution (Supplementary Fig. S5). At the later recording time points however, both the ALS groups showed an increase in connections of medium strength relative to the healthy controls. The functional connectivity was detected using the co-occurrences, and their distribution is shown in Supplementary Fig. S5. After pruning the network to remove the weakest connections, the giant component was extracted and used for further analysis. Descriptive characteristics of the size and densities of the functional networks are shown in Supplementary Fig. S6. The healthy control group contained very few networks at the first three recordings due to low network activity at this early stage in development.

Both the healthy networks and the ALS networks developed hallmarks of efficient information processing, as indicated by the high levels of local and global efficiency and Small-World Propensity (Supplementary Fig. S7). No significant differences between the groups were observed in these metrics. In correspondence with the connection strength distribution shown in Supplementary Fig. S5, the mean connection strength was higher in the untreated and TEA-treated ALS networks compared to the healthy networks at the later recording time points (Fig. 3A), but the mixed-effects model revealed no significant difference between the groups (Fig. 3B). Nonetheless, the connection strength in the TEA-treated ALS networks appeared to temporarily increase during and after the second application of TEA at 29 DIV until 34 DIV, after which it decreased and stabilized at levels comparable to that of the untreated ALS networks. The mean degree of the nodes was significantly higher in the untreated ALS networks compared to the healthy networks (p<0.01) and between the TEA-treated ALS networks and the healthy networks (p<0.05; Fig. 3C-D). There was a temporary decrease in the mean degree of the TEA-treated ALS networks during the days of perturbation and the following day, but no significant difference was observed between the two ALS groups. The mean clustering appeared to be higher in the healthy networks at the earlier time points in spite of some variations between the recordings, but decreased over time(Fig. 3E). In contrast, the clustering increased over time and was higher in the ALS groups at the later time points. Interestingly, the clustering increased in the TEA-treated ALS networks to levels comparable to the healthy networks between 29 and 34 DIV. Regardless of these observable trends, the mixed-effects model revealed no significant differences between the groups (Fig. 3F). The average path length followed a trend similar to the one observed in the mean clustering but in the opposite direction, i.e., the average path length was shorter in the healthy networks at the earlier time points, but increased over time, and was shorter in the ALS networks at the later time points (Fig. 3G). The average path length in the TEA-treated ALS networks decreased during the TEA applications compared to the untreated ALS networks and despite an increase over time, the average shortest path remained lower in the TEA-treated ALS networks until the last two recordings at 44 and 46 DIV, compared to both the untreated ALS networks and the healthy networks. While this trend was reflected in the estimated marginal means of the mixed-effects models, it was not significant (Fig. 3H). Both the ALS groups had comparable betweenness centrality across all the recordings (Fig. 3I). The betweenness centrality was however significantly higher in the healthy networks compared to both the ALS groups (Fig. 3J). In summary, the graph measures used to characterize the networks revealed topologies consistent with computational efficiency in all three groups, and there were not major differences in the information processing capacity. Notwithstanding this finding, the ALS networks appeared to be more connected, as indicated by the increased degree and trend towards higher clustering, whereas the healthy networks had a higher proportion of gatekeeper nodes, as indicated by the higher betweenness centrality, suggesting that information spreads more easily across the network in the ALS condition. Blocking of the potassium channels in ALS networks led to temporary changes in some of these metrics, but did not substantially alter the network efficiency long-term.

**Fig. 3.**
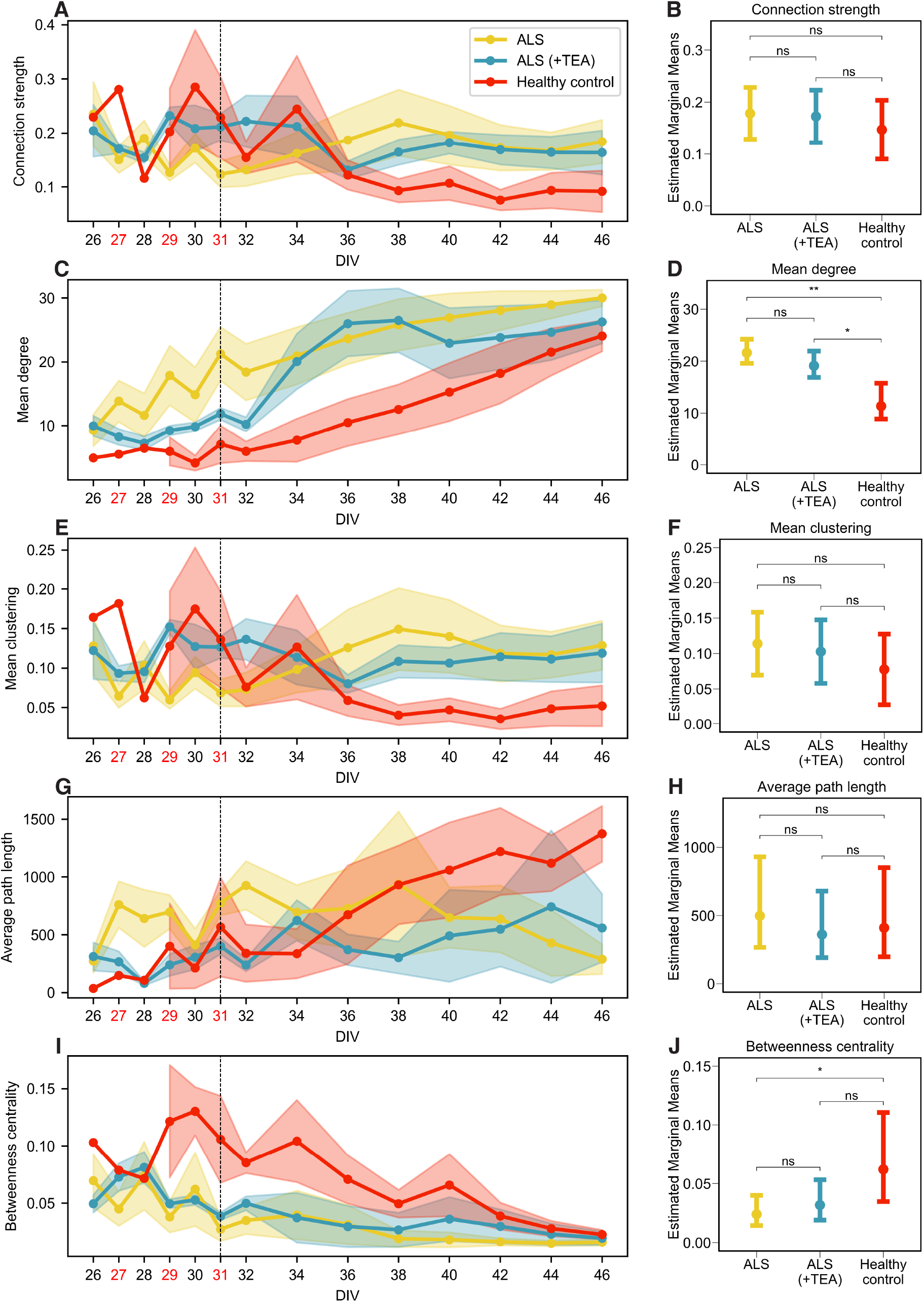
Functional connectivity and information processing capacities in motor neuron networks before and after potassium channel blocking. (**A-B**) Connection strength, represented by the mean edge weight, (**C-D**) mean degree, (**E-F**) mean clustering, (**G-H**) average shortest path length, and (**I-J**) betweenness centrality. Left column plots show the longitudinal development of the network metrics. Data is represented by the mean *±* SEM. The red-labeled days in vitro (DIV) are the days of tetraethylammonium (TEA) application, the vertical line at 31 DIV marks the final day of TEA application. Legend is shared between figures. The right column plots show mixed-effects model estimated marginal means with 95% confidence interval for each of the network metrics. * p<0.05, ** p<0.01. ns, not significant.

### Blocking of potassium channels results in upregulation of proteins involved in synaptic plasticity

To gain a better understanding of how blocking of potassium channels affected synaptic processes, we investigated the protein expression of isolated synaptosomes. We identified a total of 3050 proteins that were differentially expressed between all three groups. To investigate the overall trends in the dataset, we first conducted KEGG pathway analysis on all the DEPs. The analysis showed enrichment associated with multiple neurodegenerative diseases alongside common cellular pathways like “Metabolic pathways” and “Endocytocis” in both ALS groups compared to the healthy group, indicating a strong overlap of proteins involved in neurodegenerative pathways (Fig. 4A). We next used GO GSEA to identify the dysregulated pathways in the ALS networks, and whether these were affected as a consequence of the potassium channel blockade. GSEA of the ALS group compared to the healthy group revealed upregulation of proteins involved in pathways associated with DNA and RNA such as “RNA processing”, “DNA repair”, “ribonucleoprotein complex” and “mRNA binding”, as well as ATP activity (Fig. 4B).

**Fig. 4.**
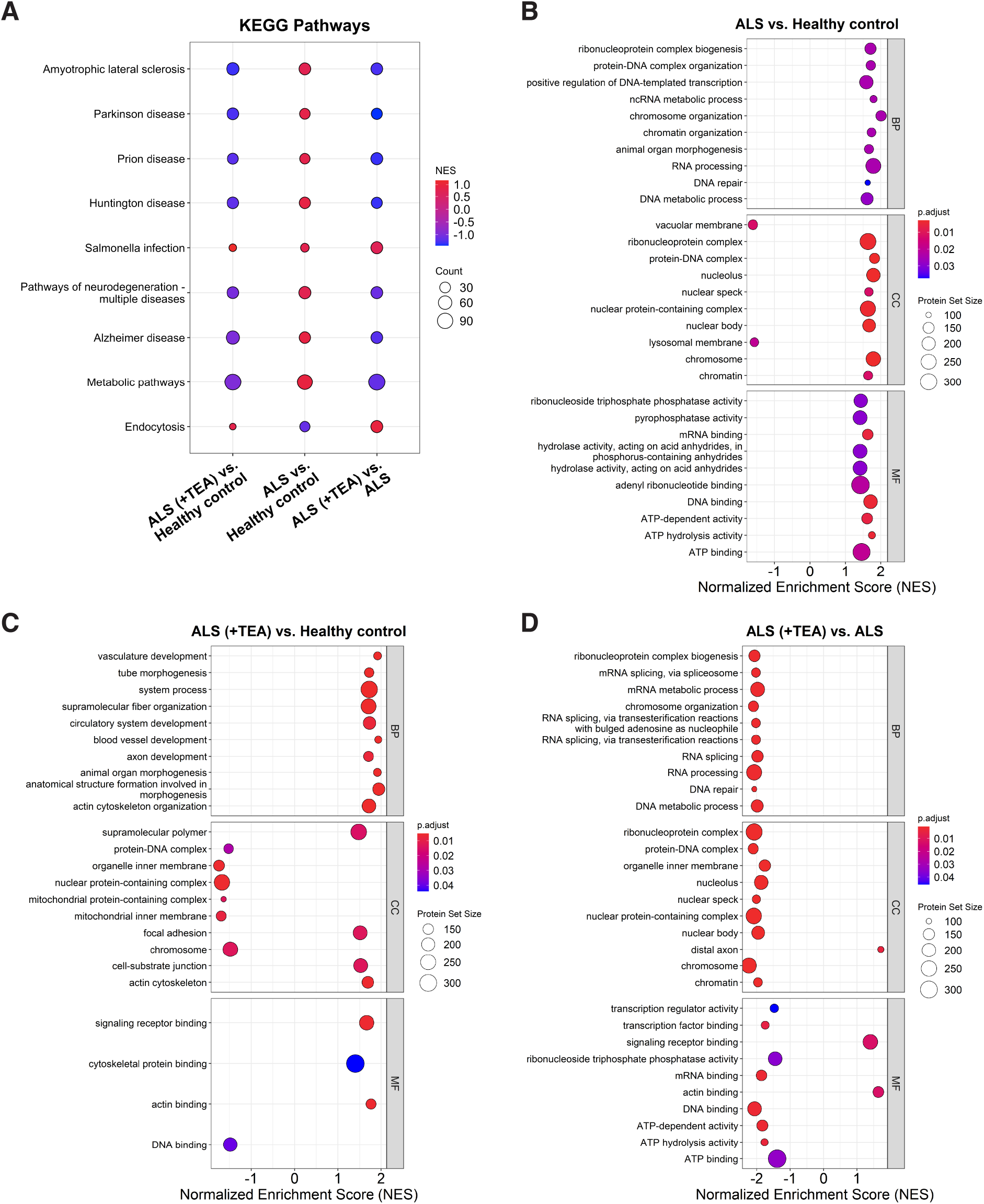
Gene set enrichment analysis revealed alterations in protein expression following temporary blocking of potassium channels in the ALS networks. (**A**) Dot plot of the top enriched Kyoto Encyclopedia of Genes and Genomes (KEGG) pathways between the three contrasts ALS (+TEA) vs. Healthy control, ALS vs. Healthy control and ALS (+TEA) vs. ALS. The color of the dots represents the Normalized Enrichment Score (NES) where positive scores indicate upregulated proteins and negative scores indicate downregulated proteins. Dot size represents the protein count. (**B**) Dot plot of the top ten enriched Gene Ontology (GO) terms for “Biological Process” (BP), “Cellular Component” (CC) and “Molecular Function” (MF) in the ALS group compared to healthy control. The x-axis represents the NES. The adjusted p-value is represented by the color of the dots, and the size of the dots represent the Protein Set Size, that is the number of proteins involved in a given GO term. (**C**) Dot plot of the top ten enriched GO terms in the TEA-treated ALS group compared to healthy control. (**D**) Dot plot of the top ten enriched GO terms in the TEA-treated ALS group compared to the untreated ALS group.

When comparing the TEA-treated ALS group to the healthy group, the majority of the upregulated proteins were associated with developmental and organizational processes, including “axon development”, “actin cytoskeleton organization” and “tube morphogenesis” (Fig. 4C). The downregulated proteins were involved in pathways associated with DNA and mitochondrial structures. To get a clearer picture of how the TEA-treated and the untreated ALS group differed, we next compared these two directly. Interestingly, the downregulated proteins in the TEA-treated ALS group were involved in many of the same pathways in which the upregulated proteins were involved in the untreated ALS group, i.e., pathways associated with DNA, RNA and ATP activity (Fig. 4D). However, the upregulated proteins were involved in “distal axon”, “signaling receptor binding” and “actin binding”.

We further investigated the differences between the TEA-treated and the untreated ALS group with GO over-representation analysis of the upregulated and downregulated proteins separately. The upregulated proteins in the TEA-treated ALS group were primarily involved in cytoskeletal organization, synaptic signaling and transmission, and neuronal structures such as “synapse” and “axon” (Fig. 5A). The downregulated proteins on the other hand, were involved in DNA- and RNA-associated processes and metabolic pathways, including “ATP-dependent activity”, “electron transfer activity” and “NADH dehydrogenase activity”, thereby corroborating the results from the GSEA (Fig. 5B). To better understand the specific processes that had been altered at the synapse in the TEA-treated ALS networks, we used the SynGO database to identify the function of the synaptic proteins. The majority of the upregulated proteins were involved in synapse organization and processes at the presynapse (Fig. 5C), while the downregulated proteins were overrepresented in metabolism and transport (Fig. 5D). A deeper investigation of the child terms associated with each of these revealed over-representation of upregulated proteins in synapse structure and assembly, synapse maturation and postsynaptic organization, as well as synaptic vesicle cycle (Fig. 5E). The downregulated proteins were primarily involved in protein translation at the pre- and postsynapse and axo-dendritic transport (Fig. 5F). Collectively, these results strongly indicated that the blocking of potassium channels engaged multiple mechanisms involved in synaptic plasticity, including reorganization of the synapse and metabolic alterations.

**Fig. 5.**
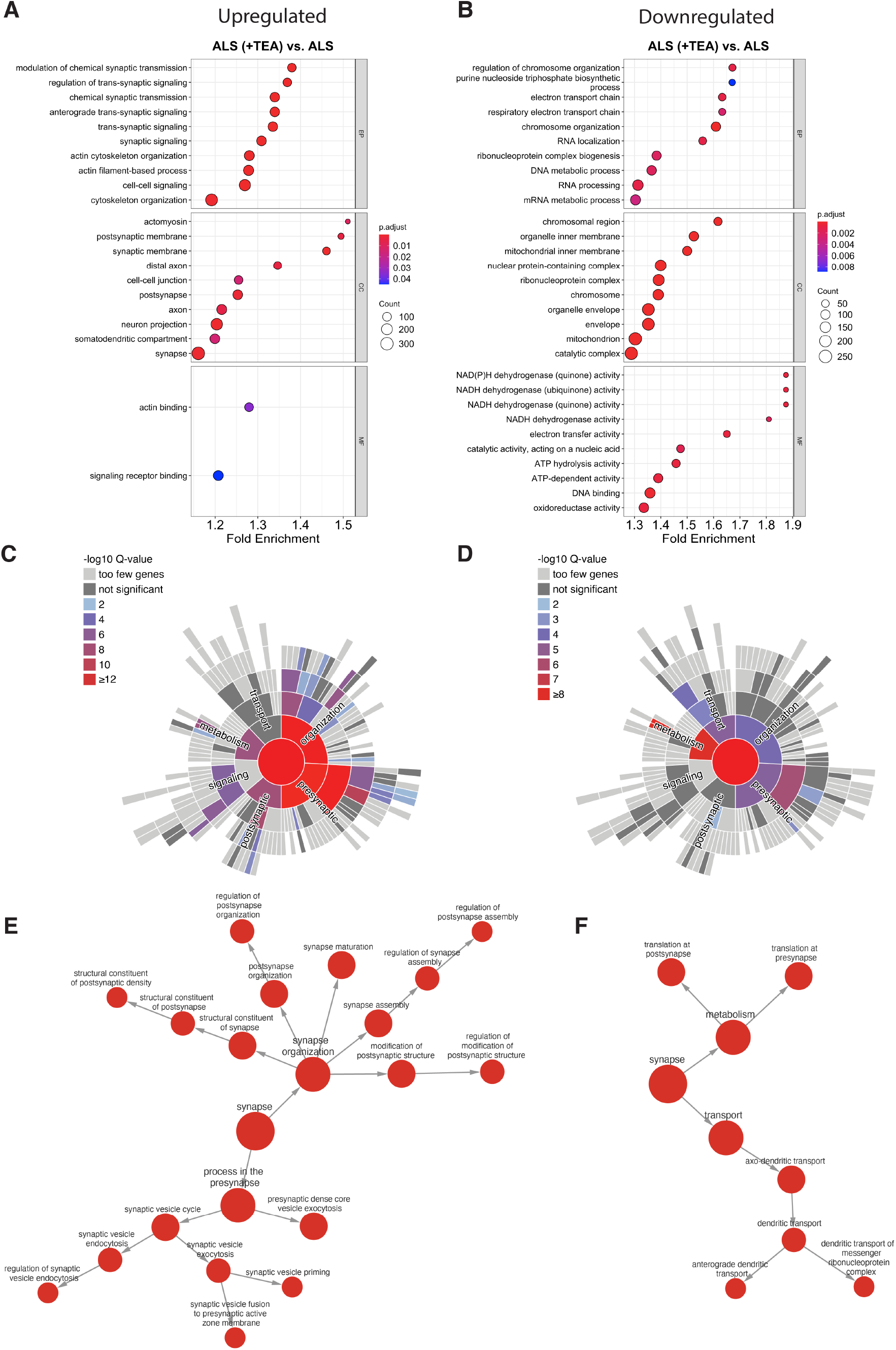
Gene Ontology (GO) over-representation analysis (ORA) revealed alterations in synaptic structure, signaling and metabolism following potassium channel blocking. (**A**) Dot plot of the top ten overrepresented GO terms for “Biological Process” (BP), “Cellular Component” (CC) and “Molecular Function” (MF) of the upregulated proteins in the tetraethylammonium (TEA)-treated ALS group relative to the untreated ALS group. The dot color represents the adjusted p-value and dot size represents the protein count associated with a specific term. (**B**) Dot plot of the top ten overrepresented GO terms of the downregulated proteins. (**C**) Enrichment of synaptic terms associated with the upregulated proteins in the TEA-treated ALS group relative to the untreated ALS group obtained from the SynGO database. (**D**) SynGO enrichment of synaptic terms associated with the downregulated proteins in the TEA-treated ALS group. (**E-F**) Networks illustrating the top two enriched synaptic terms and their associated child terms identified by the SynGO database for the upregulated (E) and downregulated proteins (F).

After investigating the coordinated changes caused by multiple proteins, we focused on the most differentially expressed proteins, that is, proteins passing the thresholds Log2 fold change≥1 and alpha=0.05. This resulted in a list of 21 proteins, and clustering of these revealed a clear separation between the healthy control group and the two ALS groups (Fig. 6A). Volcano plots were established to better visualize the difference between the ALS groups and the healthy control group (Fig. 6B-C). Interestingly, most of the 21 proteins were identified in the TEA-treated ALS group. GPR50 was the most downregulated and UTS2 the most upregulated in both the ALS groups. GPR50 is G protein-coupled receptor involved in energy metabolism and neurite outgrowth and UTS2 is a vasoconstrictor which also plays a role in neuromuscular physiology. Additionally, the signal transduction protein ARHGAP36 and ARVCF, which are involved in communication between the intra- and extracellular environments, were downregulated in both ALS groups. When comparing the TEA-treated ALS group with the healthy control group, the Ca^2+^-dependent phospholipid-binding protein CPNE1 and the voltage-gated Ca^2+^ channel CACNA1E were both downregulated. Upregulated proteins included the PDZ And LIM Domain 7 (PDLIM7) involved in cytoskeletal organization, Midkine (MDK), which promotes cell growth, migration and angiogenesis, and the membrane trafficking proteins SMAP1, WDR44 and VPS53.

**Fig. 6.**
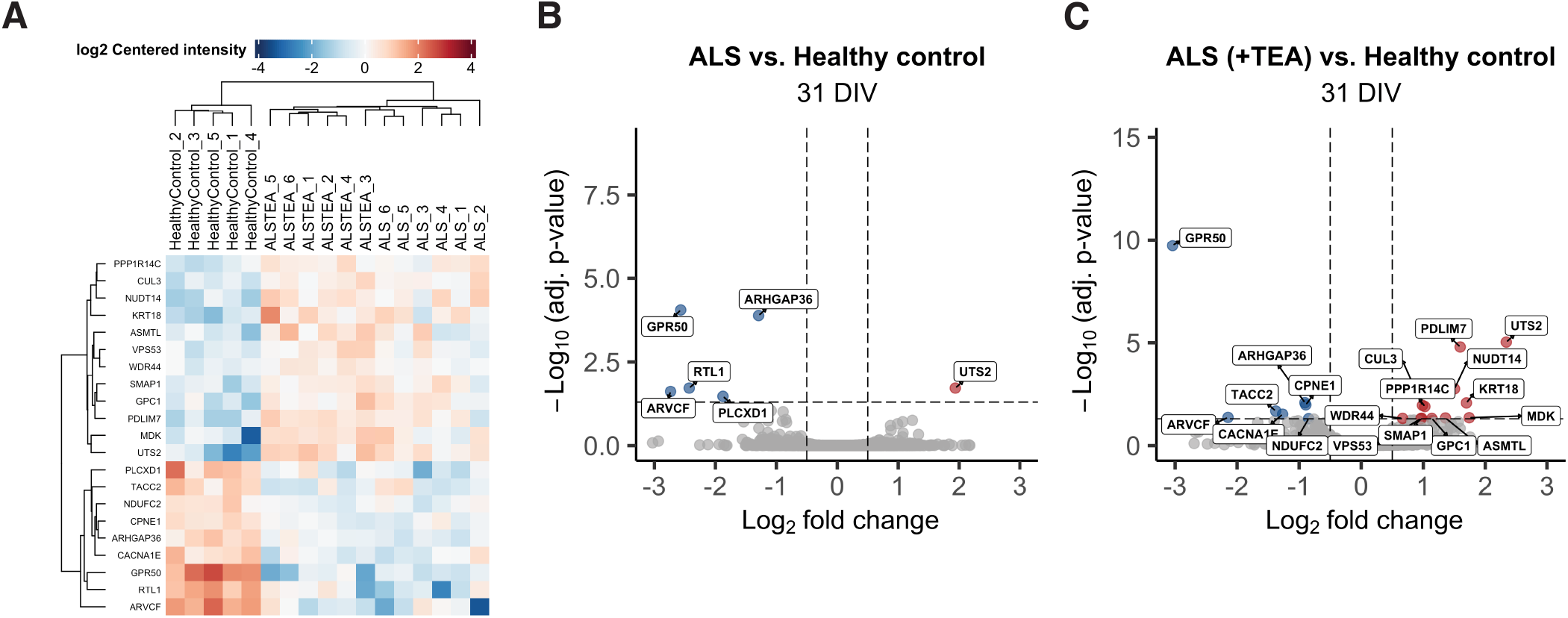
Differential protein expression analysis revealed 21 significant differentially expressed proteins (DEPs) across all three contrasts. (**A**) Heatmap of the 21 significant proteins, (**B**) Volcano plot of the upregulated (red) and downregulated (blue) significant DEPs in the ALS group compared to healthy control. Dashed lines indicate thresholds for Log2 fold change (0.5; vertical lines) and the adjusted p-value (<0.05; horizontal lines). (**C**) Volcano plot of the significant DEPs in the ALS (+TEA) group compared to healthy control.

## Discussion

This study provides novel insights into the relationship between structural alterations in ALS and the concurrent changes in neural network activity and connectivity. We show that while ALS patient-derived motor neurons develop into networks of high information processing capacity, they exhibit subtle signs of network dysfunction including increased firing rate and bursting, and increased interconnectivity. Reduced betweenness centrality further suggests that ALS patient-derived motor neurons self-organize into sub-optimal networks characterized by impaired control over the flow of network activity. Structurally, the networks exhibited increased neurite density and branching, and a time-dependent reduction in the number of dendritic spines. Additionally, proteomic analyses of motor neuron synaptosomes revealed dysregulation of metabolic pathways and protein synthesis. Importantly, by blocking the potassium channels, we observed several changes consistent with increased synaptic strength. The structural abnormalities and dysregulated molecular pathways in ALS networks were partially restored, and proteins involved in synaptic plasticity, including synapse organization and maturation, were upregulated. Furthermore, we observed reduced signs of hyperexcitability and synchronous network activity, suggesting that synaptic potentiation promoted functional network behavior associated with healthy dynamics.

The untreated ALS networks consistently exhibited shorter and more branched neurites than healthy control networks at both time points of comparison, 31 and 38 DIV (Fig. 1). This is in line with previous findings reporting aberrant neurite branching in zebrafish models with TDP-43 overexpression (74, 75) and hiPSC-based models of ALS carrying FUS mutations (76, 77). Importantly, we have previously observed aberrant neurite growth in human iPSC-derived neural network models of both ALS (25) and Parkinson’s disease (36), highlighting that altered capacity to establish and maintain synaptic connections might represent a shared deficit across different neurodegenerative diseases and associated genotypes.

Optimal wiring of synaptic connections lay the foundation for healthy network function and higher order operations of advanced neural circuits. Altered network structure and im-paired synaptic maintenance have been linked to both neurodegenerative and neurodevelopmental disorders (78, 79). We have previously shown that aberrant neurite branching and reduced neurite length in ALS patient-derived motor neurons were associated with dysregulated transcripts involved in neurite outgrowth and synaptic development and maintenance (25). Further support for the link between synaptic disruptions and altered network structure is provided by findings from cultured hippocampal neurons from C9orf72 transgenic mice (80, 81). However, these studies linked synaptic instability and dendritic spine loss to impaired neurite branching, in contrast to our findings. These differences could potentially be explained by different experiment models and morphological changes along the disease progression. In preclinical ALS models mimicking the early stages of disease, we have previously observed that compensatory mechanisms can counteract inherent disease vulnerability and microscale dysfunction, causing subtle structural and functional changes without overt signs of neurodegeneration or severe loss of computational capacity (25, 35). Hence, the increased branching observed in the present study might represent a compensatory mechanism to preserve information flow despite impaired synaptic stability and reduced long-range connectivity. Although potentially beneficial in the short term, structural rewiring is resource-demanding to uphold for longer periods of time, as indicated by the upregulation of proteins involved in energy metabolism and protein synthesis in ALS synaptosomes (Fig. 4 and Fig. 5).

Longitudinal MEA recordings revealed alterations in the network behavior of the ALS networks, specifically increased firing rate and a higher fraction of spikes in bursts and network bursts (Fig. 2). Moreover, the ALS networks were characterized by higher mean degree and increased clustering at the later time points, indicating a more interconnected network compared to the healthy controls (Fig. 3). This is consistent with a substantial body of evidence reporting hyperexcitability and hyperconnectivity in ALS patients, animal models and iPSC-derived models (2, 3, 6, 34, 82, 83). However, some studies found no clear indications of hyperexcitability (84, 85), including our own previous findings (25). Interestingly, recent evidence suggests that hyperexcitability is a transient phenotype (23, 86, 87). Prolonged overexcitation over time can lead to sodium channel inactivation and effectively block action potential firing, resulting in reduced firing, as observed in certain studies (85, 88). Furthermore, spatiotemporal network dynamics and homeostatic regulations of neuronal excitability can mask impaired firing mechanisms in individual neurons resulting in no overt signs of hyperexcitability in the collective dynamics (89). Thus, while hyperexcitability is emerging as a likely mechanism in early stages of ALS and is further corroborated by the findings in the present study, such dysfunction may not always manifest as increased firing rate at the network level.

Increased connectivity may facilitate spread of activity across large parts of the network, thereby resulting in more synchronous firing and a high proportion of spikes contained in bursts, as shown in Fig. 2C and G. Increased connectivity of functional brain networks in ALS patients has been correlated with increased synchrony (3). This also corresponds with the observed increase in neurite density and branching, but reduced average neurite length reported here, which may have been initiated to compensate for the loss of or reduced ability to establish long-range connections. A reduction in long-range connections can result in increased local connectivity and higher clustering (90), leading to more locally integrated nodes facilitating spread of activity to a large part of the network. Interestingly, we observed higher betweenness centrality in the healthy networks. Nodes with high betweenness centrality often provide the only link between other pairs of nodes and can consequently act as gate-keepers of the signaling, thereby exerting a large degree of control over the flow of information (91). The presence of these nodes may result in more functionally separated subnetworks with increased local specialization where global integration is largely controlled by the nodes with high betweenness centrality. The ALS networks on the other hand, appeared to self-organize into a suboptimal network architecture where network activity was more easily propagated globally, as indicated by the increased degree, clustering and shorter average path length. This is also supported by previous findings reporting increased connectivity and rich-club organization in brain networks following brain injury and neurodegenerative processes (35, 92, 93). Such reorganization may be initiated to preserve function in response to lost connections or inefficient signaling. However, while synchronization of network activity is necessary to maintain function, unrestrained levels of synchrony may impair the local segregation and functional specialization (94). Furthermore, maintaining such high levels of activity requires a substantial energy supply that results in rapid network failure if these needs are not met (95–97). Indeed, we identified upregulation of proteins involved in several metabolic processes in the untreated ALS synaptosomes. While increased connectivity and spread of activity may render the networks vulnerable to perturbations, the ALS networks retained properties of efficient network behavior, suggesting that these features alone are insufficient to drive pathology. However, the increased metabolic cost associated with the structural compensatory rewiring and functional reorganization may increase the vulnerability to other disease-mediating factors, thereby contributing to disease progression.

The time-dependent reduction in dendritic spine density (Fig. 1) is consistent with impaired regulation of synapses and neuronal signaling in the ALS networks. Dendritic spines are sites for advanced neural computation such as integration of synaptic signals (98, 99). Although reduced spine density did not fully distort the computational capacity of ALS networks, as shown in the electrophysiology data, increased branching and density of neurites could serve as additional potential contact points to compensate for the loss of spines. Previous work from transgenic mouse models supports the link between synaptic impairment and reduced spine density in ALS. Gorrie et al. (100) showed that hampering protein clearance by introducing an ALS-associated mutation in the UBQLN2 gene reduced dendritic spine density and led to synaptic dysfunction and cognitive deficits in mice. Impaired protein degradation and reduced spine density was also reported in a study using cultured hippocampal neurons from C9orf72 knock-out mice (81). Although these studies could suggest a loss-of-function mechanism contributing to synaptic dysfunction in ALS, other findings from cultured mouse hippocampal neurons with C9orf72 overexpression reported dendritic spine loss and synaptic impairment associated with the accumulation of toxic RNA foci and dipeptide repeat (DPR) proteins (80). While detailed investigations of RNA toxicity and DPRs were beyond the scope of this study, the proteomics data revealed upregulation of RNA processes and downregulation of lysosomal membrane mechanisms in the ALS synaptosomes (Fig. 4). This could suggest that both gain- and loss of function mechanisms contribute to the reduced dendritic spine density in the ALS networks.

The aberrant branching and reduced dendritic spine density in the ALS networks were mitigated following blocking of the potassium channels, rendering their structural characteristics more comparable to those of the healthy control networks (Fig. 1). Dendritic spines are made up of highly dynamic actin filaments, and postsynaptic densities (PSDs) are located on dendritic spines. The dendritic spine changes associated with LTP, including synthesis and maturation of spines, largely rely on actin dynamics, i.e., the polymerization and depolymerization of actin filaments by interaction with actin-binding proteins (101, 102). Consistent with this, the proteomics data revealed upregulation of actomyosin, actin and cytoskeletal organization in the TEA-treated ALS synaptosomes (Fig. 4 and Fig. 5). We also identified a presynaptic upregulation of neurotransmitter-containing secretory organelles, along with upregulation of PSD constituents and postsynaptic organization. LTP has been associated with an increased number of docked presynaptic vesicles (72, 103–105), and PSD size is positively correlated with the size of dendritic spine heads (106–108), in which an increase has been linked to LTP (72, 103–105). Since dendritic spines serve as important calcium storages, structural modifications in spines can influence calcium dynamics in neurons (72), thus linking dendritic spine dynamics and neuronal activity.

Interestingly, we did not observe an immediate increase in dendritic spines, in contrast to previous findings using the same protocol to induce LTP in rat hippocampal neurons (109). Instead, the number of spines in the TEA-treated ALS networks remained stable from 31 to 38 DIV, as opposed to the time-dependent loss of spines observed in the untreated ALS networks. Both stabilization of existing spines and the formation of new spines have been associated with LTP. For example, Xu et al. (110) reported increased formation of new spines in the mouse motor cortex after one time exposure to a paw-reaching task, and stabilization of spine number after prolonged training. Another study using photolysis of caged glutamate to induce LTP in rat hippocampal dendrites, reported increased long-term survival of spines, thereby stabilizing the synapses (111). Importantly, Stewart et al. (39) observed an increase in perforated and complex PSDs but not an overall increase in spine density following cLTP induction with TEA. Such perforation can indicate increased perimeter length of the PSD, which can enhance the size and strength of the active synaptic zone, and has been suggested to constitute a morphological correlate of increased synaptic efficacy (39, 112). We did observe an upregulation of proteins involved in synaptic organization, where the majority of proteins were involved in postsynaptic structures, including PSD (Fig. 5E). Collectively, our findings thus indicate that blocking of potassium channels improved the stabilization of dendritic spines through postsynaptic reorganization, thereby enhancing synaptic coupling and stabilizing the synapses. This may in turn have mitigated the structural deficiencies by reducing the need for compensatory branching.

Blocking of potassium channels had a clear effect on the firing rate of the motor neuron networks (Fig. 2A), which was reduced to levels comparable to that of the healthy controls on the days of TEA-application and the days immediately following. The TEA-treated ALS networks also exhibited a temporary reduction in the fraction of spikes in bursts and network bursts, and the mean burst duration. Stabilization of dendritic spines and the changes in functional network behavior appeared to facilitate a reallocation of metabolic resources, as suggested by downregulation of proteins involved in metabolic processes and protein synthesis relative to the untreated ALS networks (Fig. 4D). This indicates that reduced network activity also reduced the metabolic demand of the TEA-treated ALS networks, underscoring the relationship between excessive network activity and increased metabolic cost in ALS. Interestingly, the expression of RTL1 was modified following blocking of the potassium channels (Fig. 6). Although not commonly linked to ALS, RTL1 has been suggested to impair motor neuron innervation of skeletal muscle in Kagami-Ogata and Temple syndrome (113, 114), and regulate neuronal excitability in the locus coeruleus of mice (115). Together, these findings indicate that synaptic potentiation contributed to restore some of the biological pathways that were previously disrupted in the ALS networks.

The functional network response to TEA may appear counterintuitive as several studies have shown increased action potential firing following cLTP-induction with TEA (38–40). However, most of these studies use single-unit recordings such as voltage or patch clamp techniques to assess the effects of cLTP, and thus consider the response to induced currents as opposed to the spontaneously generated activity measured in the present study. Furthermore, we cannot rule out that the activity following cLTP induction may look different at the network level. Importantly, the mechanistic consequence of blocking potassium channels is prolongation of action potential duration by extending the repolarization phase (116–118). Longer action potential duration may strengthen the coupling between the pre- and postsynaptic neurons through increased neurotransmitter release and enhanced postsynaptic calcium influx (37). Notably, we observed an upregulation of proteins involved in synaptic vesicle cycle in the TEA-treated ALS networks compared to the untreated ALS networks, indicating increased neurotransmitter release and reuptake. The blocking of potassium channels may have resulted in longer-lasting action potentials, as opposed to more repetitive firing, thus explaining the reduced firing rate at the network level. In fact, the upregulation of proteins involved in synaptic organization and synaptic trans-mission (Fig. 5), the stabilization of dendritic spines (Fig. 1I), and the increase in functional connection strength from 29 to 34 DIV (Fig. 3A) are all consistent with synaptic potentiation and suggestive of improved synaptic stability.

Our findings clearly demonstrate several beneficial effects of synaptic strengthening in ALS patient-derived motor neuron networks. By stabilizing dendritic spines, reducing aberrant branching, and temporarily reducing excessive activity and synchrony, blocking of potassium channels promoted behavior associated with healthy network dynamics. Furthermore, several of the disrupted biological pathways in the ALS networks, including metabolic processes and protein synthesis, were stabilized following synaptic potentiation through potassium channel blocking. However, the functional dynamics did return to the levels observed in the untreated ALS networks from 34 DIV onward. This could indicate that homeostatic plasticity mechanisms were engaged to restore the global network activity back to baseline levels. This can be achieved through synaptic scaling, which modifies the synaptic strength of numerous neurons to maintain baseline average firing rates (119). Importantly, this mechanism preserves the relative differences in synaptic weights and maintains induced changes caused by strengthening at individual synapses but proportionally reduces the strength at all synapses to drive the network firing back to baseline (12). As such, synaptic scaling, likely in conjunction with other homeostatic plasticity mechanisms such as intrinsic plasticity (120), may have pushed the network activity back to a preferred baseline state after some time. Furthermore, it is possible that microscale changes in the activity of individual neurons were still present, and that such changes were not detectable at the network level.

A fundamental challenge in the search for therapeutic targets in ALS involves the progressive nature of the disease and the timing of potential interventions, which with current diagnostic tools is limited to after the onset of symptoms in patients. At this stage, the disease has already progressed far and there is a considerable pathological burden at play (121). Thus, intervention should ideally be initiated before the onset of neuronal and synaptic degeneration to stabilize function before downstream effects become devastating. Crucially, this necessitates an improved understanding of the interplay between structural and functional reconfigurations occurring early in the disease. In the present study, we demonstrate the advantages of early intervention aimed at stabilizing synaptic function. Our findings show that by interfering with fundamental mechanisms underlying neuronal excitability and action potential duration, key mechanisms underlying synaptic plasticity and LTP were promoted. The shift in structural and functional network dynamics to that of healthy networks, and upregulation of proteins involved in the organization and maturation of synapses are clear indicators of enhanced synaptic efficacy. Importantly, this contributed to restore several dysfunctions identified in the ALS networks, including increased activity, excessive synchrony and aberrant neurite branching. As such, we demonstrate a clear connection between early changes in synaptic stability and structural and functional network disruptions in ALS.

Future studies should focus on expanding this and other approaches probing synaptic plasticity to multiple patient-derived cell lines and sporadic cases. Cell lines from healthy individuals with at-risk mutations are of particular interest, as these enable the investigation of intervention even before symptom onset. Importantly, impaired plasticity in the axon initial segment have been reported to affect the regulation of intrinsic excitability in ALS hiPSC-derived motor neurons (87), and it is possible that such disruptions in plasticity mechanisms can interfere with the neurons’ ability to activate and maintain adaptive plasticity programs. Furthermore, considering that ALS is a highly heterogeneous multisystem disease, a multi-target approach may therefore be a viable future approach to promote long-term beneficial effects. Pathological mechanisms to be targeted in conjunction with synaptic strengthening may include mitochondrial dysfunction (122), neuroinflammation (123) and mutation-specific mechanisms.

In conclusion, we show that temporary blocking of potassium channels in ALS patient-derived motor neuron networks led to upregulation of proteins in synaptic organization, structure, maturation and synaptic vesicle cycle, and stabilization of dendritic spines, consistent with enhanced synaptic potentiation and LTP. Stabilization of the synapses may in turn have led to reduced aberrant neurite branching and density, and promoted functional network dynamics consistent with healthy behavior, including reduced signs of hyperactivity and excessive synchrony. Taken together, the findings of this study emphasize the role of early synaptic impairment in ALS and demonstrate the potential of engaging fundamental aspects of synaptic plasticity to restore network function in neurodegenerative diseases.

## AUTHOR CONTRIBUTIONS

**AMK:** Conceptualization, Methodology, Software, Formal Analysis, Investigation, Data Curation, Writing - Original Draft, Writing - Review & Editing, Visualization. **MBG:** Methodology, Software, Formal Analysis, Investigation, Writing - Original Draft, Writing - Review & Editing, Visualization. **NC:** Methodology, Software, Formal Analysis, Investigation, Writing - Review & Editing, Visualization. **AS:** Conceptualization, Resources, Writing - Review & Editing, Funding acquisition. **IS:** Conceptualization, Resources, Writing - Review & Editing, Supervision, Funding acquisition.

## ACKNOWLEDGEMENTS

Mass spectrometry-based analyses were performed by the Proteomics and Modomics Experimental Core (PROMEC), Norwegian University of Science and Technology (NTNU) and The Central Norway Regional Health Authority. This facility is a member of the National Network of Advanced Proteomics Infrastructure (NAPI), which is funded by the Research Council of Norway INFRASTRUKTUR-program (project number: 295910). Data storage and handling is supported under the NIRD/Notur project NN9036K. Confocal imaging was performed by Marianne Sandvold Beckwith at the Cellular and Molecular Imaging Core Facility (CMIC), Norwegian University of Science and Technology (NTNU). CMIC is funded by the Faculty of Medicine and Health Sciences at NTNU and Central Norway Regional Health Authority.

## SUPPLEMENTARY MATERIALS

The Supplementary Materials file contains:

Fig. S1-S7

Table S1-S3

