## Supplemental Materials for "Induced long-term potentiation improves synaptic stability and restores network function in ALS motor neurons"

Figure S1-S7
Table S1-S3

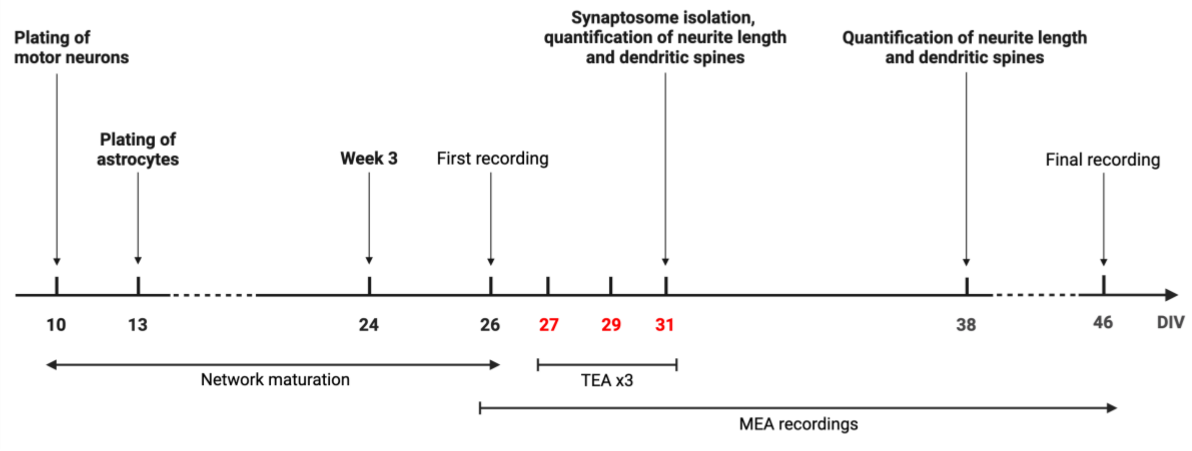

Figure S1. Experimental timeline. Human induced pluripotent stem cells (hiPSCs) were differentiated into motor neurons, and plated onto micro-electrode arrays (MEAs), 6-well plates and 8-well chamber slides at 10 days in vitro (DIV). Astrocytes were seeded at 13 DIV. The first MEA recording was conducted at 26 DIV, and the networks were subsequently recorded every other day until 46 DIV. Chemical long-term potentiation (cLTP) was induced using tetraethylammonium (TEA) in half of the ALS networks at 27, 29 and 31 DIV, followed by washout and recording of network activity to assess the immediate effects of the perturbation. At 31 DIV, immediately following the final TEA application, synaptosomes were isolated for protein expression assays and networks were fixed for immunocytochemistry assessing neurite length and dendritic spines. One week later, at 38 DIV, additional immunocytochemistry assays were conducted to quantify neurite length and dendritic spines. Figure created with BioRender.com.

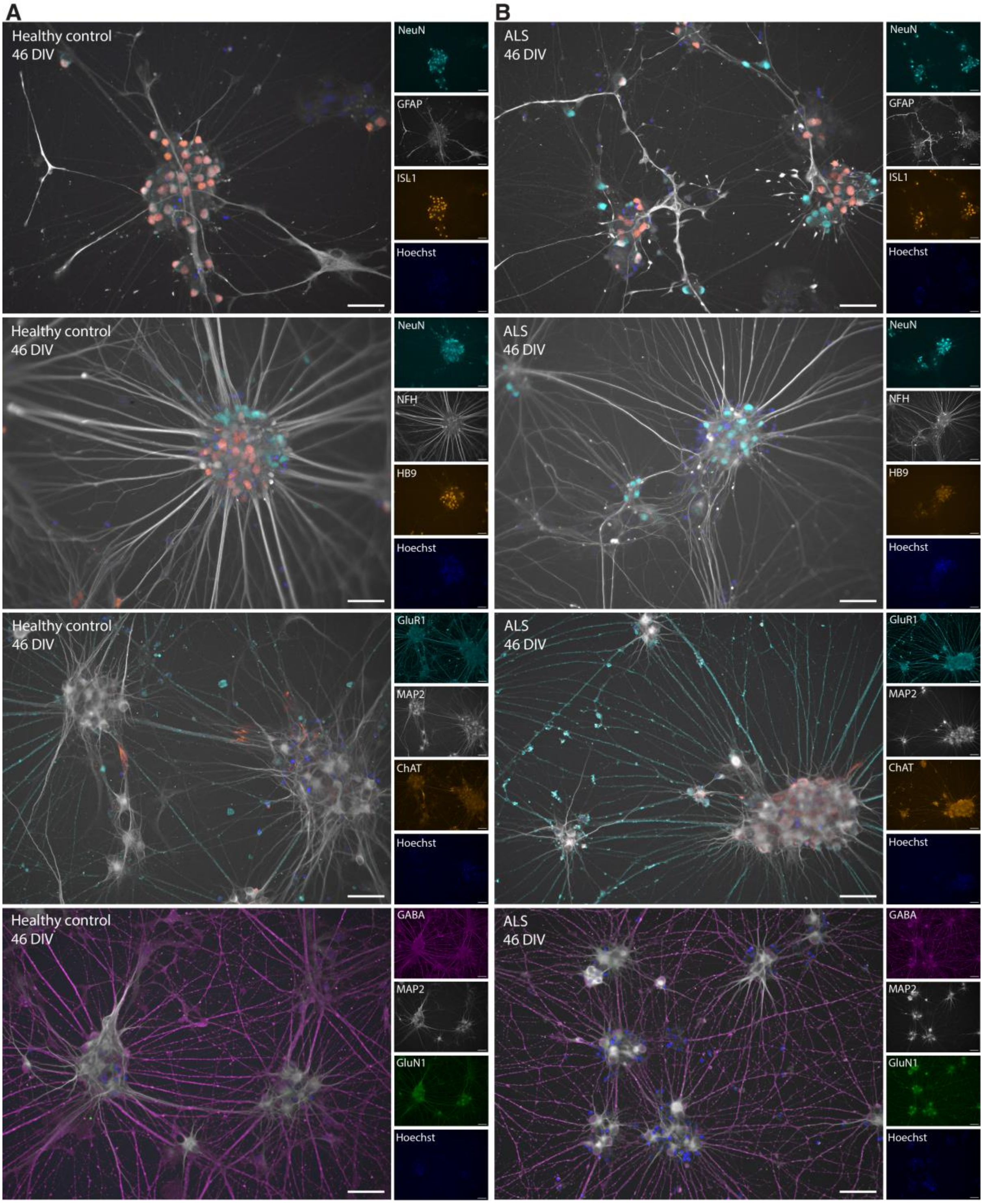

Figure S2. Immunocytochemistry assays confirming the expression of motor neuron markers and receptors. (**A**) Healthy motor neurons and (**B**) ALS patient-specific motor neurons both expressed the motor neuron markers ISL1, HB9 and ChAT at 46 DIV. Positive expression of NeuN, NFH and MAP2 indicated maturity of the networks with a high degree of clustering and axonal bundling, and GFAP confirmed the presence of astrocytes. The AMPA-receptor subunit GluR1 and the NMDA-receptor subunit GluN1 were also expressed, as well as the inhibitory neurotransmitter GABA. Scale bars: 100 μm.

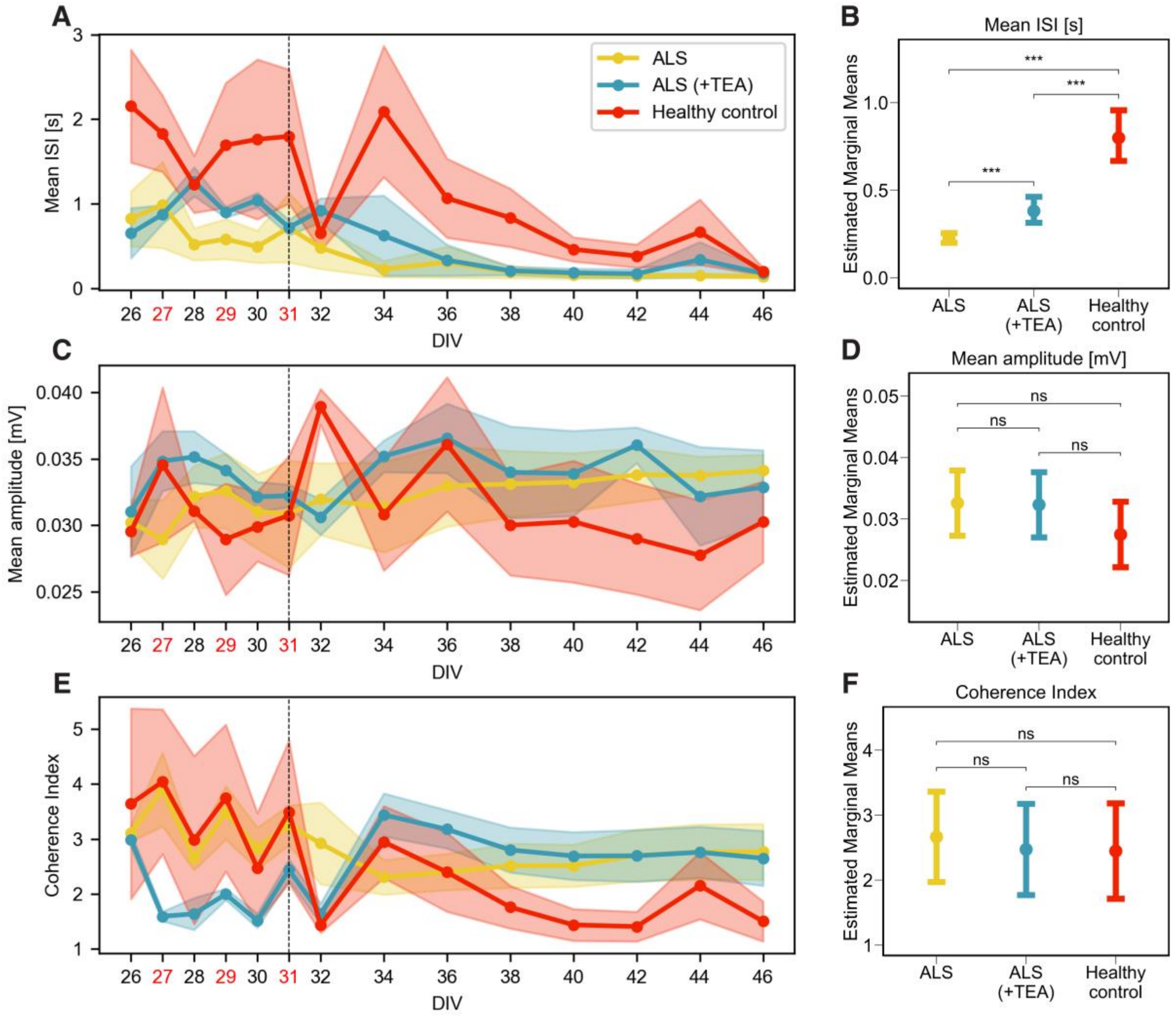

Figure S3. Additional network activity parameters characterizing the functional development of motor neuron networks. (**A-B**) Mean inter-spike interval (ISI) in seconds, (**C-D**) mean amplitude in mV, and (**E-F**) Coherence Index representing the network synchrony. Left column plots: Functional development over time. Data is represented by mean values for all networks in each group (solid lines and circles). Shaded area shows the SEM. The red-labeled days in vitro (DIV) are the days of tetraethylammonium (TEA) application, the vertical line at 31 DIV marks the final day of TEA application. Legend shared between figures. Right column plots: Mixed-effects model estimated marginal means with 95% confidence interval for each of the network parameters. * $p<0.05$, ** $p<0.01$, *** $p<0.001$. ns, not significant.

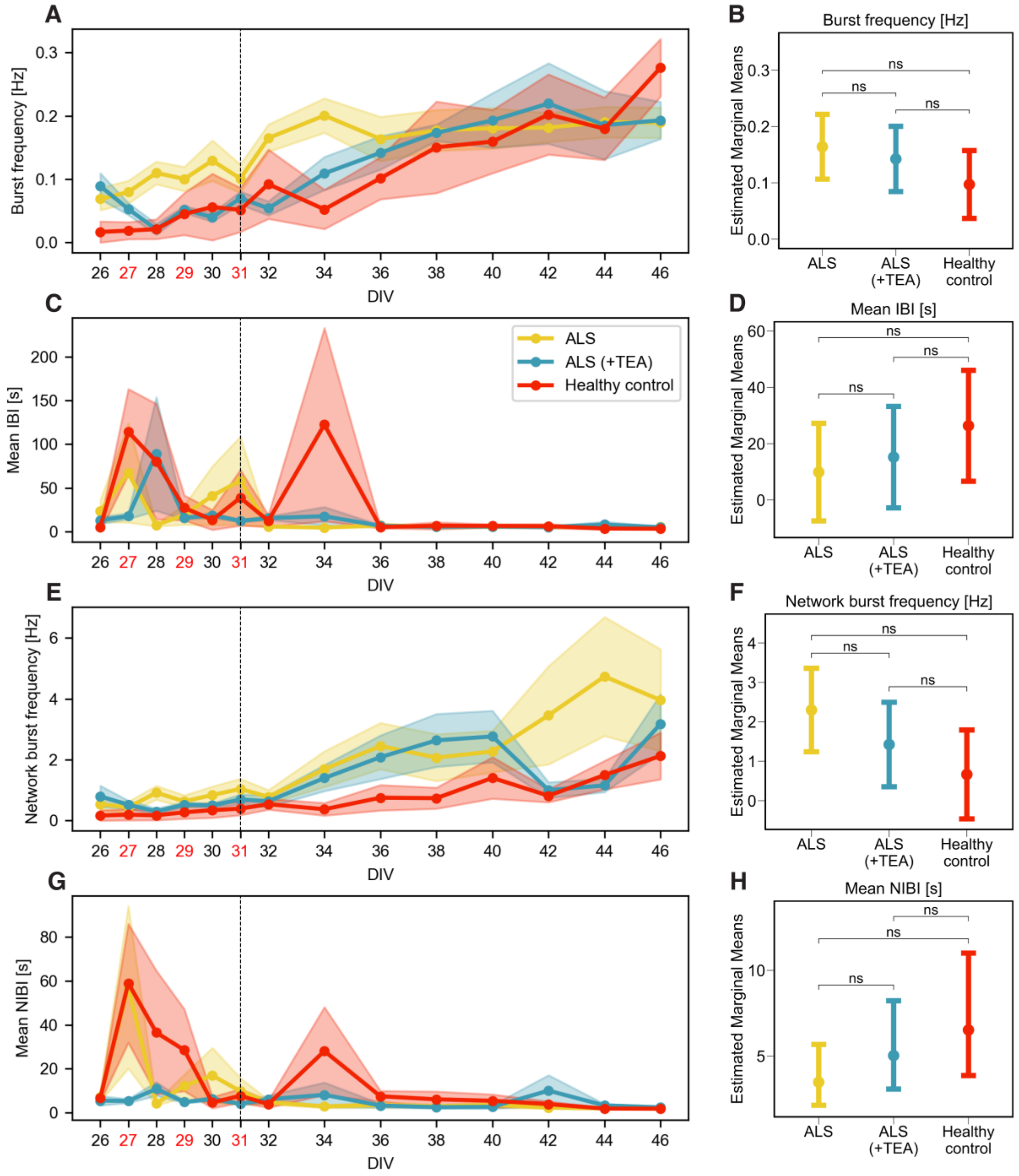

Figure S4. Characterization of bursting properties before and after blocking of potassium channels. (**A-B**) Burst frequency in Hz, (**C-D**) mean inter-burst interval (IBI) in seconds, (**E-F**) network burst frequency in Hz, and (**G-H**) mean network inter-burst interval (NIBI) in seconds. Left column plots: Development of bursting characteristics over time. Data is represented by the mean ± SEM. Legend shared between figures. Right column plots show the mixed-effects model estimated marginal means with 95% confidence interval for each of the network parameters. * $p<0.05$, ** $p<0.01$, *** $p<0.001$. ns, not significant.

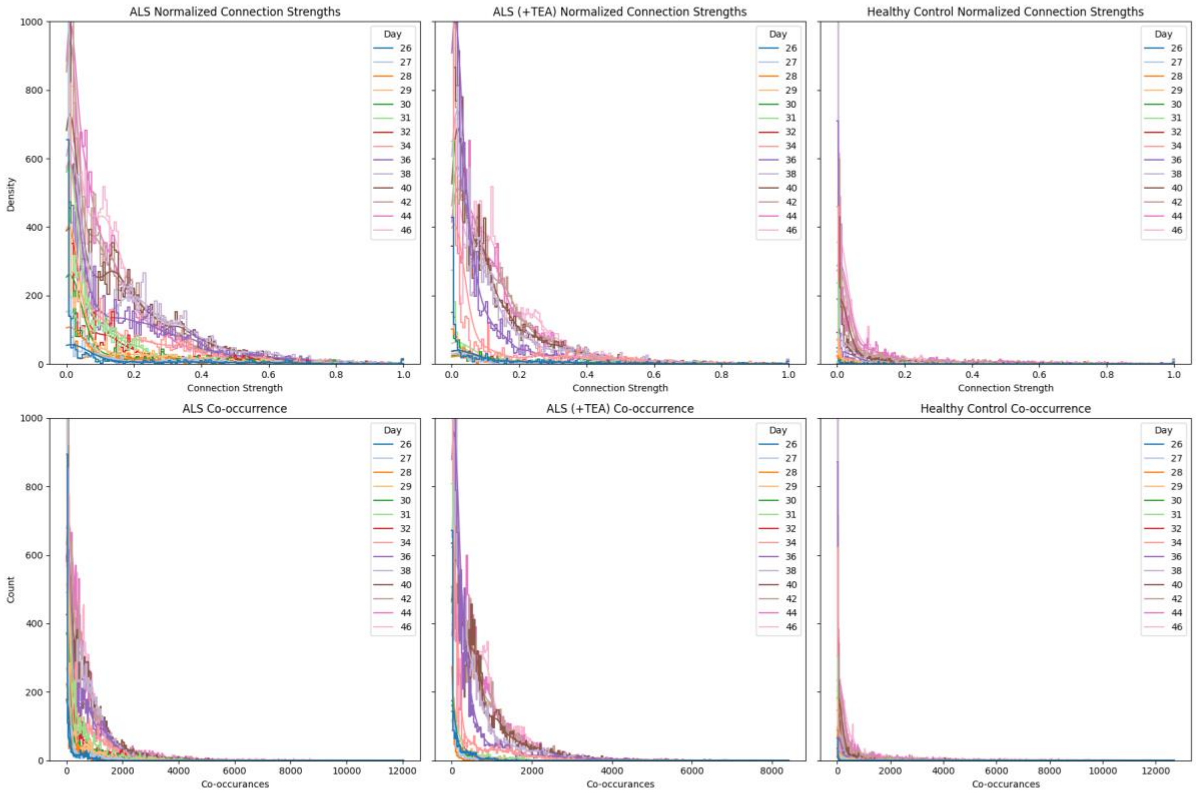

Figure S5. Distributions of the normalized connection strengths and co-occurrances identified in the ALS networks (left column), the TEA-treated ALS networks (middle column) and in the healthy control networks (right column). Top panels show the distributions of the weights with corresponding kernel density estimate of all identified connections pooled across all wells at each recording time point. Bottom panels show the distributions of all identified co-occurrances pooled across all wells at each time point.

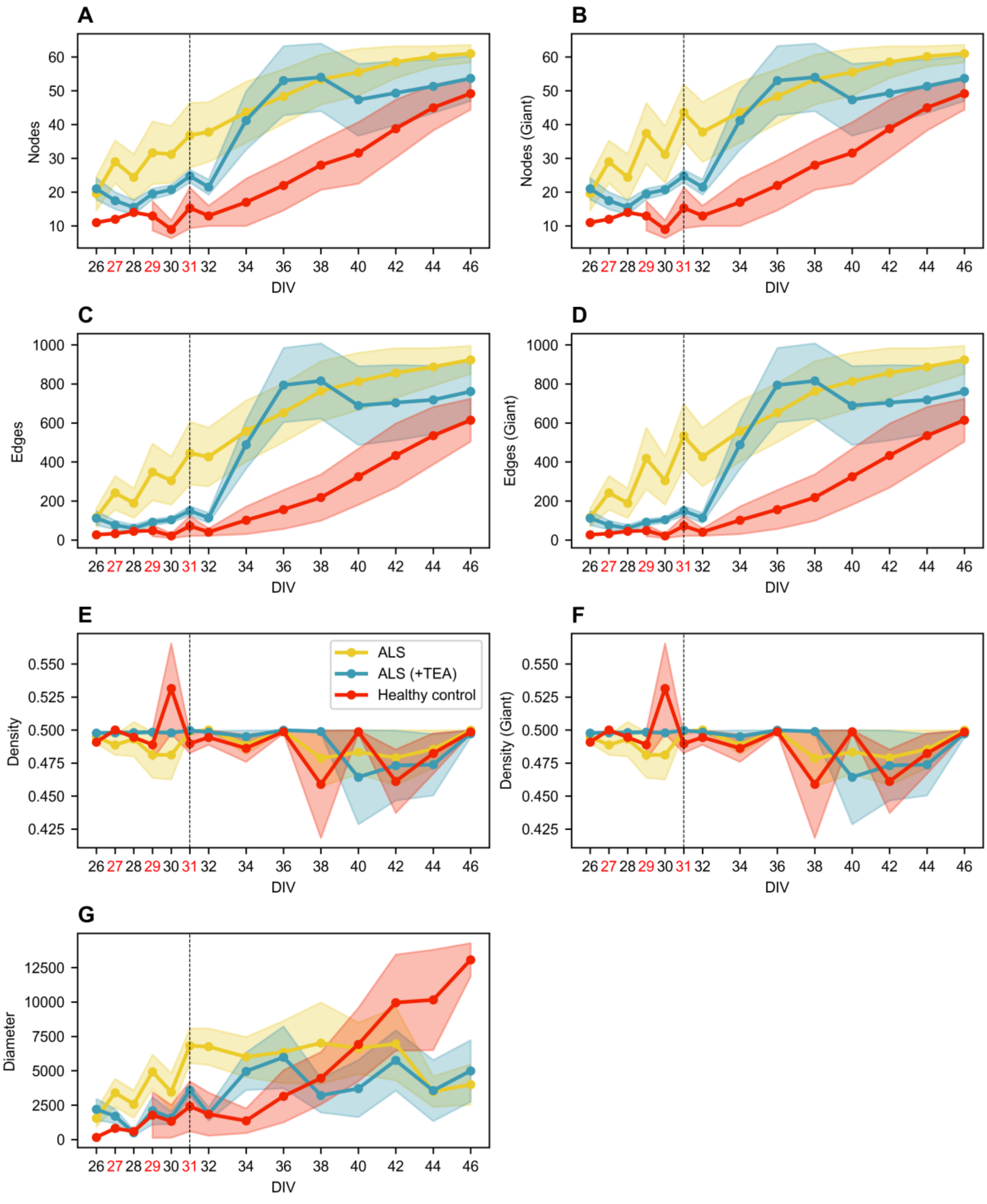

Figure S6. Descriptive characteristics of the networks used in the functional connectivity analysis. (**A**) Number of nodes, (**B**) number of nodes in the giant component, (**C**) number of edges, (**D**) number of edges in the giant component, (**E**) network density, (**F**) network density in the giant component, (**G**) network diameter calculated on the inverse weights of the giant component. Data is represented by the mean ± SEM. Legend shared between figures.

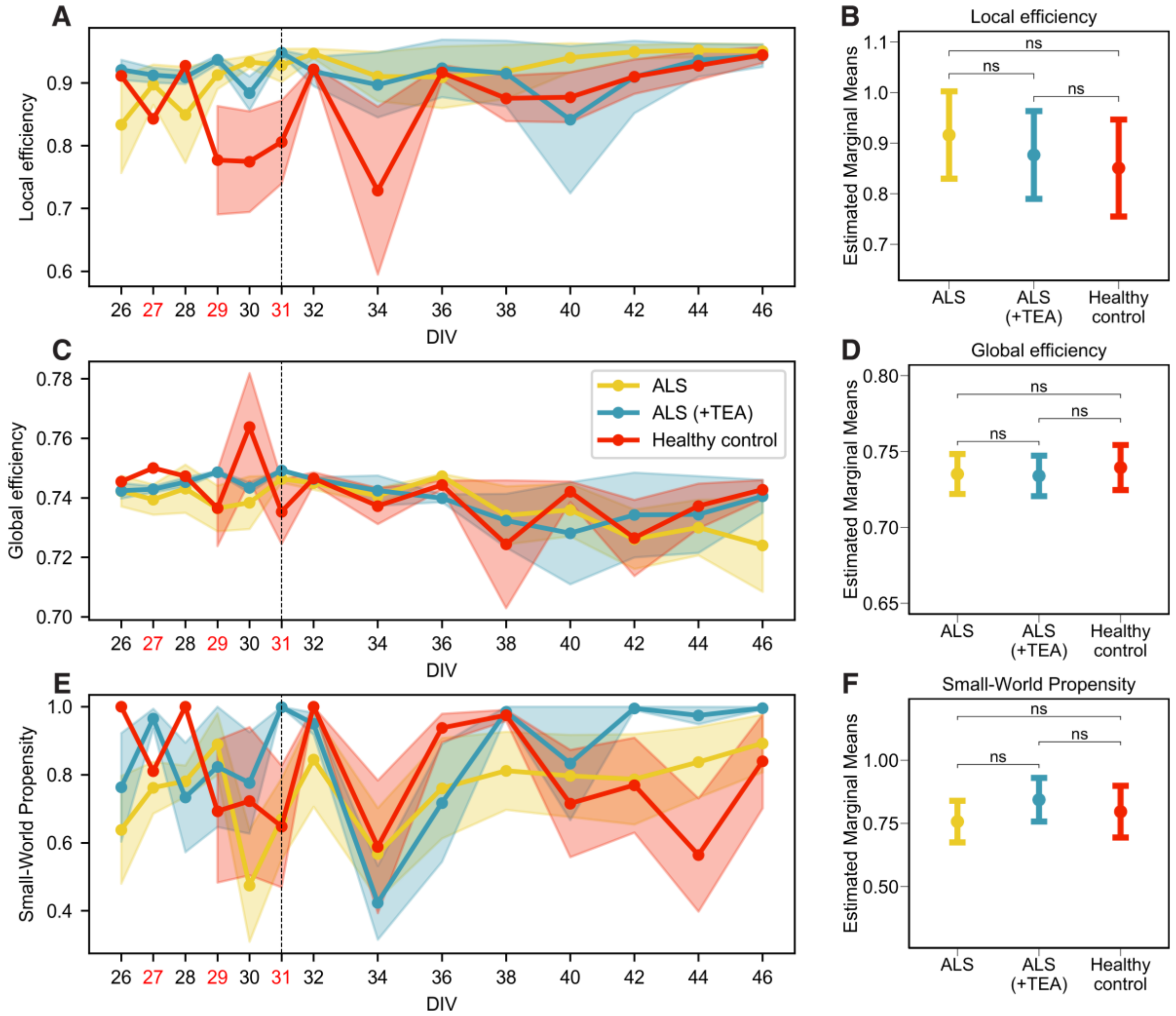

Figure S7. Efficiency metrics and Small-World Propensity. (**A-B**) Local efficiency, (**C-D**) Global efficiency, (**E-F**) Small-World Propensity, where values above 0.6 are considered small-world. The development over time is shown in the left-column plots. Data is represented by the mean ± SEM. Legend shared between figures. Right column plots show the mixed-effects model estimated marginal means with 95% confidence interval for each of the network parameters. ns, not significant.

Table S1. Overview of the total number of neural networks included in the network activity analysis at each individual day in vitro (DIV).

| **DIV** | **26** | **27** | **28** | **29** | **30** | **31** | **32** | **34** | **36** | **38** | **40** | **42** | **44** | **46** |
|---|---|---|---|---|---|---|---|---|---|---|---|---|---|---|
| ALS | 5 | 6 | 5 | 6 | 6 | 6 | 6 | 6 | 6 | 6 | 6 | 6 | 6 | 6 |
| ALS (+TEA) | 4 | 4 | 4 | 4 | 4 | 4 | 4 | 5 | 5 | 6 | 6 | 6 | 6 | 6 |
| Healthy control | 2 | 3 | 2 | 3 | 3 | 4 | 2 | 5 | 5 | 5 | 5 | 5 | 6 | 5 |

Table S2. Overview of the total number of neural networks included in the functional connectivity analysis at each individual day in vitro (DIV).

| **DIV** | **26** | **27** | **28** | **29** | **30** | **31** | **32** | **34** | **36** | **38** | **40** | **42** | **44** | **46** |
|---|---|---|---|---|---|---|---|---|---|---|---|---|---|---|
| ALS | 5 | 5 | 5 | 6 | 5 | 6 | 5 | 6 | 6 | 6 | 6 | 6 | 6 | 6 |
| ALS (+TEA) | 4 | 4 | 4 | 4 | 4 | 4 | 4 | 5 | 5 | 5 | 6 | 6 | 6 | 6 |
| Healthy control | 1 | 1 | 1 | 3 | 3 | 3 | 2 | 4 | 4 | 4 | 5 | 5 | 5 | 5 |

Table S3. Results from mixed-effects models applied to each network parameter. The model and distribution fitted for each parameter is shown under the "Mixed-effects model" column with the link function in parentheses. Models with a log or inverse link function transform the response variable, and were back-transformed to the original response scale in the plots shown in Figure 2, 3, S3, S4 and S7. GLMM, generalized linear mixed-effects model; LMM, linear mixed-effects model; TEA, tetraethylammonium; EMM, Estimated Marginal Means; SE, standard error of the mean; CI, confidence interval.

| *Network parameter* | *Mixed-effects model* | *Condition* | *EMM* | *SE* | *Lower CI* | *Upper CI* |
|---|---|---|---|---|---|---|
| Firing rate [Hz] | GLMM gaussian (log) | ALS | 1.72 | 0.200 | 1.333 | 2.12 |
| | | ALS (+TEA) | 1.40 | 0.204 | 1.003 | 1.80 |
| | | Healthy control | 0.68 | 0.233 | 0.224 | 1.14 |
| Mean ISI [s] | GLMM inverse gaussian (log) | ALS | -1.491 | 0.0629 | -1.614 | -1.3675 |
| | | ALS (+TEA) | -0.965 | 0.0987 | -1.159 | -0.7717 |
| | | Healthy control | -0.225 | 0.0920 | -0.405 | -0.0443 |
| Mean amplitude [mV] | GLMM inverse gaussian (identity) | ALS | 0.0326 | 0.00271 | 0.0273 | 0.0379 |
| | | ALS (+TEA) | 0.0323 | 0.00271 | 0.0270 | 0.0376 |
| | | Healthy control | 0.0275 | 0.00272 | 0.0222 | 0.0328 |
| Coherence Index | LMM gaussian (identity) | ALS | 2.66 | 0.324 | 1.97 | 3.36 |
| | | ALS (+TEA) | 2.47 | 0.328 | 1.77 | 3.17 |
| | | Healthy control | 2.45 | 0.347 | 1.71 | 3.18 |
| Fraction of spikes in bursts | LMM gaussian (identity) | ALS | 0.819 | 0.0865 | 0.634 | 1.00 |
| | | ALS (+TEA) | 0.653 | 0.0872 | 0.467 | 0.84 |
| | | Healthy control | 0.337 | 0.0908 | 0.145 | 0.53 |
| Mean burst duration [s] | LMM gaussian (identity) | ALS | 0.406 | 0.0438 | 0.3121 | 0.500 |
| | | ALS (+TEA) | 0.358 | 0.0441 | 0.2638 | 0.452 |
| | | Healthy control | 0.176 | 0.0457 | 0.0789 | 0.273 |
| Fraction of spikes in network bursts | LMM gaussian (identity) | ALS | 0.830 | 0.0913 | 0.634 | 1.027 |
| | | ALS (+TEA) | 0.695 | 0.0924 | 0.498 | 0.893 |
| | | Healthy control | 0.420 | 0.0971 | 0.214 | 0.626 |
| Mean network burst duration [s] | GLMM gamma (log) | ALS | -1.53 | 0.193 | -1.91 | -1.154 |
| | | ALS (+TEA) | -1.30 | 0.198 | -1.69 | -0.912 |
| | | Healthy control | -1.21 | 0.214 | -1.63 | -0.793 |

| | | | | | | |
|---|---|---|---|---|---|---|
| Burst frequency [Hz] | LMM gaussian (identity) | ALS | 0.1641 | 0.0268 | 0.1066 | 0.222 |
| | | ALS (+TEA) | 0.1424 | 0.0271 | 0.0845 | 0.200 |
| | | Healthy control | 0.0971 | 0.0284 | 0.0369 | 0.157 |
| Mean IBI [s] | LMM gaussian (identity) | ALS | 9.89 | 7.97 | -7.43 | 27.2 |
| | | ALS (+TEA) | 15.22 | 8.39 | -2.81 | 33.2 |
| | | Healthy control | 26.36 | 9.31 | 6.66 | 46.1 |
| Network burst frequency [Hz] | LMM gaussian (identity) | ALS | 2.300 | 0.493 | 1.241 | 3.36 |
| | | ALS (+TEA) | 1.425 | 0.501 | 0.354 | 2.50 |
| | | Healthy control | 0.666 | 0.532 | -0.461 | 1.79 |
| Mean NIBI [s] | GLMM gamma (log) | ALS | 1.25 | 0.250 | 0.757 | 1.74 |
| | | ALS (+TEA) | 1.62 | 0.251 | 1.123 | 2.11 |
| | | Healthy control | 1.88 | 0.267 | 1.352 | 2.40 |
| Connection strength | LMM gaussian (identity) | ALS | 0.178 | 0.0230 | 0.1285 | 0.228 |
| | | ALS (+TEA) | 0.172 | 0.0236 | 0.1218 | 0.223 |
| | | Healthy control | 0.147 | 0.0267 | 0.0903 | 0.204 |
| Mean degree | GLMM gaussian (inverse) | ALS | 0.0462 | 0.00252 | 0.0412 | 0.0511 |
| | | ALS (+TEA) | 0.0524 | 0.00350 | 0.0455 | 0.0592 |
| | | Healthy control | 0.0880 | 0.01250 | 0.0635 | 0.1125 |
| Mean clustering | LMM gaussian (identity) | ALS | 0.1138 | 0.0206 | 0.0695 | 0.158 |
| | | ALS (+TEA) | 0.1027 | 0.0209 | 0.0578 | 0.147 |
| | | Healthy control | 0.0775 | 0.0234 | 0.0277 | 0.127 |
| Average path length | GLMM gamma (log) | ALS | 6.21 | 0.319 | 5.59 | 6.84 |
| | | ALS (+TEA) | 5.89 | 0.323 | 5.26 | 6.52 |
| | | Healthy control | 6.02 | 0.373 | 5.29 | 6.75 |
| Betweenness centrality | GLMM inverse gaussian (log) | ALS | -3.73 | 0.261 | -4.24 | -3.22 |
| | | ALS (+TEA) | -3.45 | 0.263 | -3.96 | -2.93 |
| | | Healthy control | -2.78 | 0.295 | -3.36 | -2.20 |
| Local efficiency | LMM gaussian (identity) | ALS | 0.916 | 0.0401 | 0.830 | 1.002 |
| | | ALS (+TEA) | 0.877 | 0.0405 | 0.790 | 0.963 |
| | | Healthy control | 0.851 | 0.0448 | 0.755 | 0.947 |

| | | | | | | |
|---|---|---|---|---|---|---|
| Global efficiency | LMM gaussian (identity) | ALS | 0.735 | 0.00613 | 0.722 | 0.748 |
| | | ALS (+TEA) | 0.734 | 0.00623 | 0.721 | 0.747 |
| | | Healthy control | 0.739 | 0.00698 | 0.725 | 0.754 |
| Small-World Propensity | LMM gaussian (identity) | ALS | 0.757 | 0.0375 | 0.675 | 0.839 |
| | | ALS (+TEA) | 0.843 | 0.0397 | 0.757 | 0.930 |
| | | Healthy control | 0.796 | 0.0484 | 0.694 | 0.899 |
